# Condensates of disordered proteins have small-world network structures and interfaces defined by expanded conformations

**DOI:** 10.1101/2022.05.21.492916

**Authors:** Mina Farag, Samuel R. Cohen, Wade M. Borcherds, Anne Bremer, Tanja Mittag, Rohit V. Pappu

## Abstract

The formation of membraneless biomolecular condensates is driven by macromolecules with sticker-and-spacer architectures that undergo phase separation coupled to percolation (PSCP). Driving forces for PSCP are governed by the interplay between reversible inter-sticker crosslinks and solvation preferences of spacers. Here, we introduce molecular and mesoscale descriptions of structures within, outside, and at the interfaces of condensates that are formed by prion-like low complexity domains (PLCDs), which are exemplars of intrinsically disordered, linear multivalent proteins. Our studies are based on simulations that accurately describe sequence-specific phase behaviors of PLCDs. We find that networks of reversible, intermolecular, inter-sticker crosslinks organize PLCDs into small-world topologies within condensates. These topologies result from distinct conformational preferences within dense, dilute, and interfacial regions. Specifically, the degree of conformational expansion varies non-monotonically, being most expanded at the interface and most compact in the dilute phase with molecules preferring to be oriented perpendicular to condensate interfaces. This contrasts with dense and dilute phases where molecules are randomly oriented relative to one another. Our results demonstrate that even simple condensates, with only one type of macromolecule, feature inhomogeneous spatial organizations of molecules and interfacial features that likely prime them for being locations of biochemical activity.

---

In living cells, many proteins and nucleic acids are concentrated into membraneless biomolecular condensates that form and disassemble at the right place and time ^1, 2, 3, 4^. Macromolecular phase separation has emerged as the dominant theme for explaining how condensates form and dissolve in response to environmental, mechanical, chemical, and developmental cues ^5^. Multivalence of domains or motifs that form reversible physical crosslinks are defining features of proteins that are biologically relevant drivers of condensate formation ^1, 6, 7^. Recent attention has focused on proteins with disordered regions known as prion-like low complexity domains (PLCDs) ^8^. The compositional makeup of PLCD sequences is distinctive. Roughly 60-70% are polar residues, 15-20% of the residues are aromatic π-systems, and the remainder are ionizable residues ^9^. Within each PLCD, the aromatic residues are distributed uniformly across the linear sequence ^8, 10, 11^. Although sequences of PLCDs vary considerably across evolution, the compositional biases and linear patterning of aromatic residues are conserved features ^10, 12^. Recent experimental work has uncovered the physiochemical principles underlying the connections between sequence-encoded features and the driving forces for phase separation of PLCDs. These studies used the PLCD from the protein hnRNPA1, hereafter referred to as the A1-LCD, and designed variants thereof as targets for investigation ^8, 9^.

Here, we build on the extant knowledge base regarding phase behaviors of A1-LCD and designed variants of this system to investigate the molecular and mesoscale organization of proteins within, outside, and at the interfaces of condensates. This work is motivated by the realization that PLCDs and other proteins that are drivers of bio-macromolecular phase separation are biological instantiations of linear associative polymers ^7^. Specifically, intrinsically disordered, linear multivalent proteins that drive phase separation are instantiations of linear associative polymers ^7, 13, 14, 15, 16, 17, 18, 19, 20, 21^. Such systems are defined by sticker-and-spacer architectures, whereby the driving forces for phase separation are governed by the interplay between physical crosslinks among stickers, and the effective, solvent-mediated interactions among spacers ^7, 20, 22^. Accordingly, the phase behaviors of sticker-and-spacer systems are characterized as phase separation coupled to percolation (PSCP) ^6, 7, 20, 21, 22, 23^. This process generates condensates that are microgel-like ^24, 25^, implying that the physically crosslinked networks of molecules are condensate spanning ^7, 20, 22^. Such systems are viscoelastic in nature and their material properties are governed by emergent structures of the underlying networks, including the conformations of individual molecules, the extent of crosslinking they enable, the topological structures generated by crosslinking, and the impacts of spacers on the dynamics of intermolecular rearrangements that drive the making and breaking of crosslinks ^26, 27^.

Here, we undertake a systematic investigation of molecular and mesoscale structural descriptions of condensate interiors, interfaces, and coexisting dilute phases. For this, we leverage residue-level descriptions afforded by simulations that use LaSSI, a bond fluctuation-based lattice model paradigm ^20^, designed to reproduce the macroscopic phase behavior of the A1-LCD system and numerous designed variants of this system ^9^. Our analysis of structural properties of condensate interiors, interfaces, and coexisting dilute phases yields insights into complexities that are manifest even for condensates formed by seemingly simple systems such as PLCDs with sticker-and-spacer architectures.

## Computational sticker-and-spacer model for A1-LCD and designed variants

We used LaSSI, which is a lattice-based simulation engine for coarse-grained simulations of sequence- and / or architecture-specific PSCP of biopolymers. The development of LaSSI was inspired by the bond fluctuation model for lattice polymers ^28, 29^. Specifically, LaSSI is a generalization of the bond fluctuation model developed by Shaffer ^30^. In the current implementation, we use a single bead-per-residue version of the LaSSI model. There are nine specific residue types, one each for tyrosine (Y), phenylalanine (F), arginine (R), lysine (K), glycine (G), serine (S), threonine (T), glutamine (Q), asparagine (N), and a generic residue (X). The contact energies between pairs of sites occupied by the different residue types were parameterized using a protocol described in the Methods section and summarized in Extended Data Fig. 1.

Size exclusion chromatography-aided small-angle x-ray scattering (SEC-SAXS) data were collected for the A1-LCD and a series of designed variants ^9^. These data provide an estimate of the ensemble-averaged radius of gyration (*R*_*g*_) for each of the PLCDs at 25°C while ensuring that proteins do not undergo phase separation or oligomerization ^9^. We developed a model for the contact energies among all unique pairs of residue types using the following protocol: We performed simulations of individual chains, computed the correlation between LaSSI-derived and measured chain dimensions, and iterated to convergence via a Gaussian process Bayesian optimization approach developed in previous work ^31^. The resultant model for the contact energies is summarized in Fig. 1A. In dimensionless units, the optimized pairwise contact energies range from ≈ -20 to ≈ -0.4.

**Fig 1:**
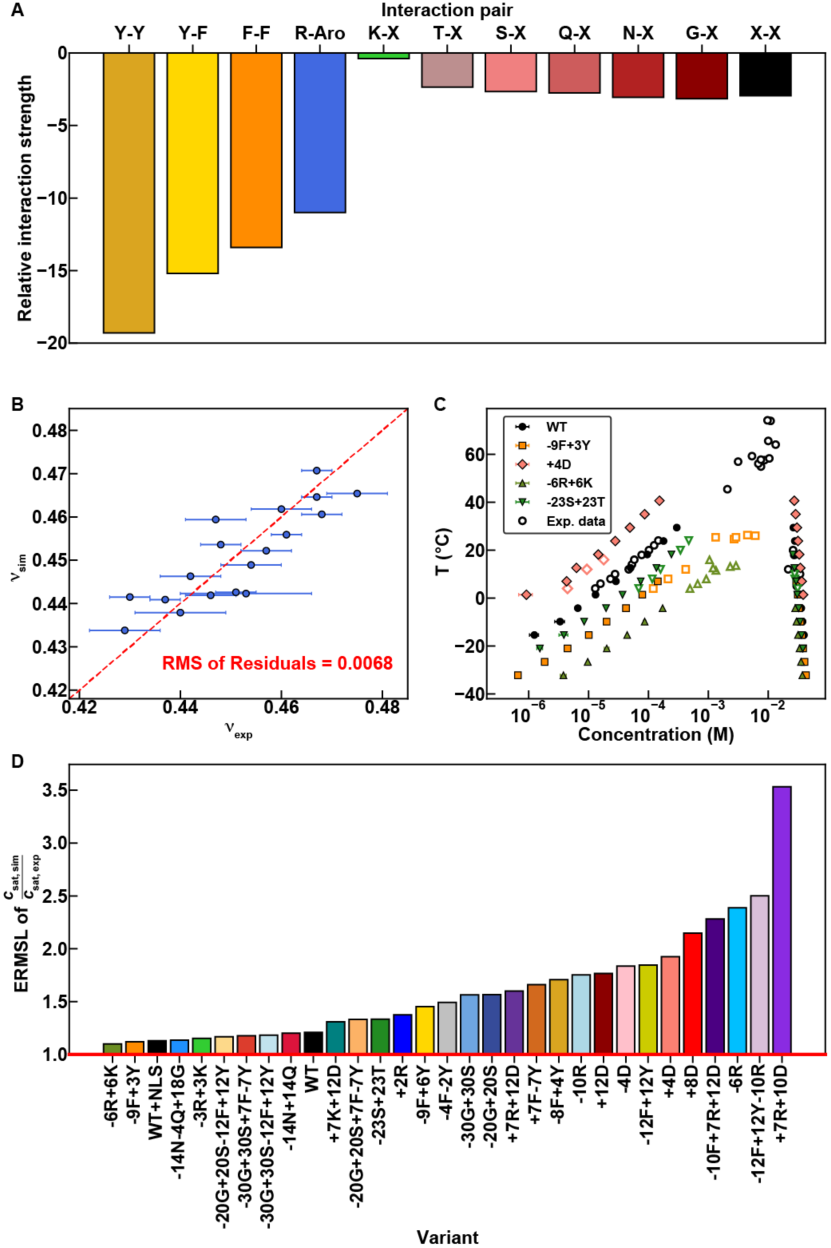
Setup and assessment of the computational model. **(A)** Pairwise interaction strengths used in the computational model. Amino acids are referred to by single-letter codes. “X” is used to indicate any amino acid for which a specific interaction is not defined. “Aro” is used to indicate either tyrosine or phenylalanine. Contact energies for Y-Y, Y-F, F-F, R-Aro, and X-X were parameterized using Gaussian process Bayesian optimization (GPBO; see Methods and Extended Data Figure 1). All other energies were parameterized by matching experimental and computational phase diagrams of “spacer” variants ^9^. **(B)** *R*_*g*_ values scale with chain length according to the relation *R*_*g*_ ∼ *N*^ν^. Here, ν is an apparent scaling exponent ν_app_, that is sequence specific, and is extracted from SEC-SAXS data using the approach developed by Riback et al., ^32^. We compare values of ν obtained by fitting SEC-SAXS data to a molecular form factor (ν_exp_) to those obtained from single-chain LaSSI simulations (ν_sim_) and use GPBO to parameterize a computational model. Each data point corresponds to a unique A1-LCD variant. The red dashed line represents the regime where ν_exp_ = ν_sim_, and the root mean squared error is calculated using the residuals from this line. Error bars represent standard errors derived from the fit to the molecular form factor. **(C)** Calculated phase diagrams (solid markers) of various A1-LCD variants plotted alongside experimental phase diagrams (open markers). Temperature and concentration are converted from simulation units to experimental units using the same scaling factors for each variant. These factors were prescribed by Martin et al., ^8^. Error bars represent standard errors from the mean across 3 replicates. **(D)** ERMSL (see Methods) comparing experimentally measured (*c*_sat, exp_) and computationally derived (*c*_sat, sim_) saturation concentrations.

We use Metropolis Monte Carlo simulations to sample configurational space for single chains and multiple chains on a cubic lattice ^20^. Accordingly, the transition probability for converting between pairs of configurations is proportional to exp(-Δ*E*/*k*_*B*_*T*). Here, Δ*E* is the difference in energy between a pair of configurations. In the simulations, we set *k*_B_ = 1, and *T* is in the interval 40 ≤ *T* ≤ 60. In units of the dimensionless simulation temperature, replacing Y-Y interactions with a Y-K interaction, which represents the largest change in Δ*E*, will range from ≈ 0.32 *k*_*B*_*T* to 0.47 *k*_*B*_*T*, depending on the simulation temperature. That the model reproduces the target function against which it was parameterized is evident in Fig. 1B, which shows a strong positive correlation between the apparent scaling exponents inferred from SEC-SAXS measurements and from the LaSSI simulations of individual chain molecules.

Note that our parameterization of the model rests on the assumption of a strong coupling between the driving forces for single-chain compaction and phase separation ^8, 33, 34^. Bremer et al., showed that this coupling breaks down for variants where the net charge per residue (NCPR) deviates from zero in a way that does not impact single chain dimensions, but does impact multi-chain interactions ^9^. Based on the analysis of Bremer et al.,^9^ we included a mean-field NCPR-based adjustment to the potentials for simulations of multichain phase behavior. In these simulations, the pairwise interactions were weakened or strengthened by an amount that is proportional to the difference in NCPR values between that of the given variant and that of the wild type (see Methods and Extended Data Fig. 2).

## Judging the accuracy of computed phase diagrams

We computed coexistence curves (binodals) for 31 different sequences including the wild-type A1-LCD (Extended Data Fig. 3). Results for the wild-type and four other variants studied by Bremer et al.,^9^ are shown in Fig. 1C. Simulation temperatures were converted to degree-Celsius and volume fractions were converted to molar units using the conversion factors introduced by Martin et al., ^8^. The computed and measured binodals show good agreement with one another. For each of the 31 sequences, we calculated the exponential root mean square log (ERMSL) between the measured and computed low concentration arms of binodals (see Methods). The ERMSL is a positive value greater than or equal to 1. An ERMSL value of 10 indicates that, on average, the concentrations along the low concentration arm of a binodal differ by an order of magnitude from the measured values. Alternatively, an ERMSL value of 1 indicates that there is no error between the dilute arms and that they should overlay perfectly. For all but one of the sequences, the ERMSL is ≤ 2.5 (Fig. 1D). This shows that the model reproduces measured phase boundaries for all experimentally characterized variants even though we parameterized the model using SEC-SAXS data for only 50% of the sequences.

## Conformations in dense phases are more expanded compared to the coexisting dilute phases

We quantified the *R*_*g*_ values of individual chain molecules in coexisting dilute and dense phases of our simulations. The results are shown in Fig. 2A for the wild-type A1-LCD. Here, *R*_*g*_ is plotted against the parameter 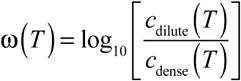, which is the temperature-dependent width of the two-phase regime (Extended Data Fig. 4A). Note that ω is negative because the concentration in the dilute phase (*c*_dilute_) is lower than the concentration in the dense phase (*c*_dense_). Also note that ω increases with *T* and approaches zero as *T* approaches the critical temperature *T*_*c*_ ≈ 49°C beyond which the system exits from the two-phase regime. In both the dilute and dense phases, the *R*_*g*_ values of individual molecules increase as *T* increases (Fig. 2A). However, for each of the temperatures that are below *T*_*c*_, the *R*_*g*_ values in the dense phase are systematically higher than *R*_*g*_ values in the dilute phase (Fig. 2A). This is due to the network of intermolecular interactions that are realized in the dense phase as opposed to the intramolecular interactions in the dilute phase – a feature that is depicted pictorially in Fig. 2B.

**Fig. 2:**
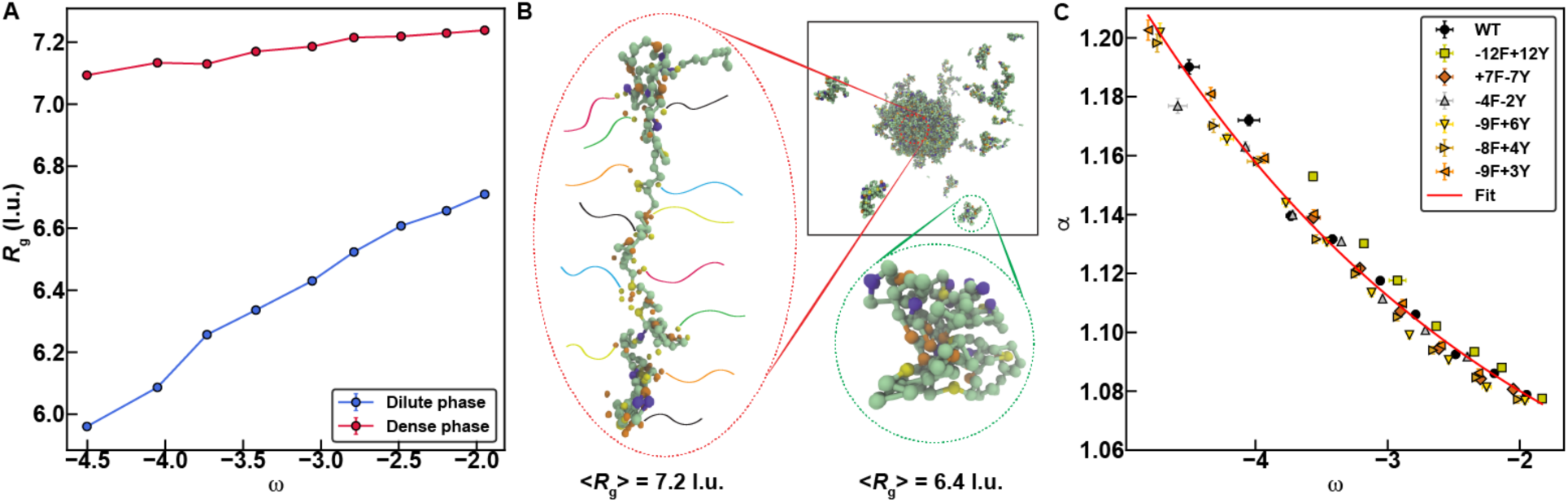
Comparison of conformations within dense vs. dilute phases. **(A)** *R*_*g*_ of chains in the dilute (blue) and dense phase (red) derived from condensates of the wild-type A1-LCD plotted against the width of the two-phase regime (see text). **(B)** A schematic depicting how intramolecular sticker-sticker interactions promote chain compaction in the dilute phase, whereas intermolecular sticker-sticker interactions promote chain expansion in the dense phase. **(C)** Swelling ratio, which quantifies the degree of expansion of chains in the dense phase relative to the dilute phase, is plotted against the width of the two-phase regime for specific A1-LCD variants. The datapoints collapse onto a single exponential curve (solid red curve; see Methods for fitting model and parameters). Error bars represent standard errors across 3 replicates. l.u. is lattice units.

Recently, Hazra and Levy showed that generic polymers featuring a mixture of long- and short-range interactions are relatively more expanded in dense vs. coexisting dilute phases ^35^ – a result we had reported prior to their publication ^36^. Given two distinct observations of similar phenomena using very different models, we analyzed results for variants where we either titrated the number of aromatic stickers or we altered the identities of the aromatic stickers Y vs. F. The goal was to assess the robustness of chain swelling across the phase boundary. For this, we computed the swelling ratio α, defined as the ratio of *R*_*g*_ in the dense phase to *R*_*g*_ in the dilute phase. We note that α approaches unity as *T* tends to *T*_c_ (Extended Data Fig. 4B). As with A1-LCD, we find that the mutational variants are more expanded in the dense phase when compared to the dilute phase (Fig. 2C). Interestingly, in a plot of α against ω (Fig. 2C), we find that the swelling ratios for seven distinct variants collapse onto a single master curve without any adjustable parameters. This curve can be fit to an exponential decay function (Fig. 2C). This implies that knowledge of the width of the two-phase regime for a disordered PLCD allows us to infer the swelling ratio from the master curve. Further, if we supplement knowledge regarding the width of the two-phase regime with measurements of chain dimensions in the dilute phase, then we can use a master curve to infer the average *R*_*g*_ values of individual chain molecules in the dense phase, at least for PLCDs.

We analyzed the three-way interplay of intra-chain, inter-chain, and chain-solvent contacts as determinants of *R*_*g*_ in the dense phase (Extended Data Fig. 5). Here, chain-solvent contacts refer to the observation of a vacant site adjacent to a site occupied by a chain. Our analysis shows that the sole determinant of the extent of chain compaction is the fraction of intramolecular contacts (*f*_intra_) (Extended Data Fig. 5). For a given *R*_g_ value, which fixes *f*_intra_, the sum of the fractions of inter-chain (*f*_inter_) and chain-solvent contacts (*f*_sol_) is constrained by: *f*_intra_ + *f*_inter_ + *f*_sol_ = 1. Accordingly, *f*_inter_ + *f*_sol_ = (1 – *f*_intra_), and hence any increase in *f*_sol_ is compensated by a decrease in *f*_inter_ and vice versa (Extended Data Fig. 5).

## Networking of chains within dense phases is determined by the strengths and valence of stickers

From the contact energies (ε) summarized in Fig. 1A we note that the interaction strengths of stickers follow a hierarchy whereby ε_YY_ > ε_YF_ > ε_FF_ > ε_RY/F_. Therefore, it follows that tyrosine (Y), and phenylalanine (F) are the primary stickers whereas arginine (R) is an auxiliary sticker in PLCDs. Stickers form reversible crosslinks, and in the lattice simulations a crosslink is distinguished from a random contact by the frequency of observing a specific pair of residues coming into contact. Crosslinking is governed by the hierarchy of interaction energies and the temperature, specifically the distance from the critical point. We quantified a ratio of association *g*_*a*_, which we define as 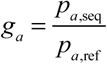. Here, *p*_*a*,seq_ is the relative probability of observing sticker-sticker vs. sticker-spacer contacts in the sequence (seq) of interest. The parameter *p*_*a*,ref_ is the homopolymer equivalent of *p*_*a*,seq_. The homopolymer is of the same length as the wild-type A1-LCD. The contact energies, which are identical among all residues, are parameterized to reproduce the computed binodals for the wild-type A1-LCD. For comparative analysis, we impose the sticker-and-spacer architecture of the wild-type sequence onto the homopolymer (Extended Data Fig. 6).

The ratios of association were computed for each of the 31 sequence variants (Extended Data Fig. 7A-C). The ratio of association is largest in the variant where all phenylalanine residues are replaced by tyrosine (see data for -12F+12Y in Extended Data Fig. 7A). Replacing all tyrosine residues with phenylalanine lowers the ratio of association (see data for +7F-7Y in Extended Data Fig. 7A). Decreasing the valence of aromatic residues, whereby six of the stickers in A1-LCD are replaced by spacers, causes a lowering of the ratio of association to be below one. This implies that the extent of networking is weakened even when compared to the equivalent homopolymer (see data for -4F-2Y in Extended Data Fig. 7A). Surprisingly, replacing auxiliary stickers such as arginine with a spacer that weakens the driving forces for phase separation increases the ratios of association when compared to the wild-type A1-LCD (see data for -3R+3K and -6R+6K compared to the wild-type A1-LCD in Extended Data Fig. 7A; also see panel E in Extended Data Fig. 3). This is because the auxiliary stickers compete with the primary aromatic stickers. However, even though the ratio of association of stickers is higher in variants with fewer arginine residues, the driving forces for phase separation are weakened by the competing effects of spacers with a higher preference to be solvated. In general, changes to the identities and hence interactions mediated by spacers have a negligible effect on the ratios of association as shown in our results for thirteen different variants where the sticker identities and valence are those of wild-type A1-LCD, but the identities and hence interactions mediated by spacers have been altered substantially (Extended Data Fig. 7B-C). When compared to data for measured and computed binodals (see Extended Data Fig. 3), we conclude that solubility determining interactions involving spacers can impact the driving forces for phase separation without affecting the networking of stickers. Taken together, these results demonstrate that some sequence features may affect driving forces for phase separation and internal condensate organization in non-equivalent ways. From a protein engineering standpoint, this feature could enable so-called separation of function mutations.

Next, we quantified the probability *P*(*s*) of realizing clusters of lattice sites within condensates with *s* stickers that form via inter-sticker crosslinks. Although the distributions (shown in Extended Data Fig. 7D for the wild-type A1-LCD) are exponentially bounded for small *s*, they have heavy tails. This feature also appears in the probability density for self-avoiding walks ^37^, with the difference being that the heavy tails here are created by the crosslinking stickers. We fit the data for *P*(*s*) to the functional form for the cumulative distribution function of a discrete Weibull distribution ^38^ given by:

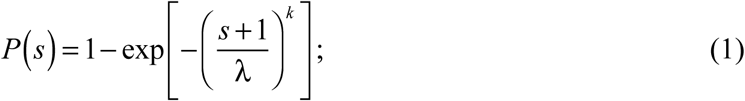

Here, *s* is the number of stickers within each cluster, whereas λ and *k* are, respectively, sequence-specific scale and shape parameters of the Weibull distribution. The sequence-specific values of λ and *k* were extracted by linear regression analysis of plots of ln[–ln(1–*P*(*s*))] vs. ln(*s*+1). As shown in Extended Date Fig. 7E, increasing the strength of stickers (−12F+12Y) leads to increased clustering of stickers (larger λ-values) when compared to wild-type A1-LCD. Likewise, decreasing the strengths of stickers (+7F-7Y) lowers the extent of clustering of stickers (lower λ-values) when compared to the wild-type A1-LCD. Lowering the valence of stickers significantly lowers the extent of clustering (see data for -4F-2Y in Extended Date Fig. 7E). Finally, the extent to which large clusters of stickers are formed, quantified by the values of *k*, where lower values imply heavier tails, is governed almost exclusively by the valence of stickers.

## Condensates form small-world structures defined by networks of physical crosslinks

The heavy-tailed nature of the cluster distributions suggests that molecules can be networked to be condensate spanning. This would generate specific types of network structures, which we analyzed using graph-theoretic methods ^39^. In this analysis of the simulation results, we treat each molecule within a condensate as a node. An undirected edge is drawn between a pair of nodes if at least one pair of stickers from the molecules in question forms a contact. The resultant graphs depicting the representative topological structures at a given snapshot are shown for the wild-type A1-LCD (panels A-C of Fig. 3). Each node is colored by its degree, which is defined as the number of edges emanating from the node.

**Fig. 3:**
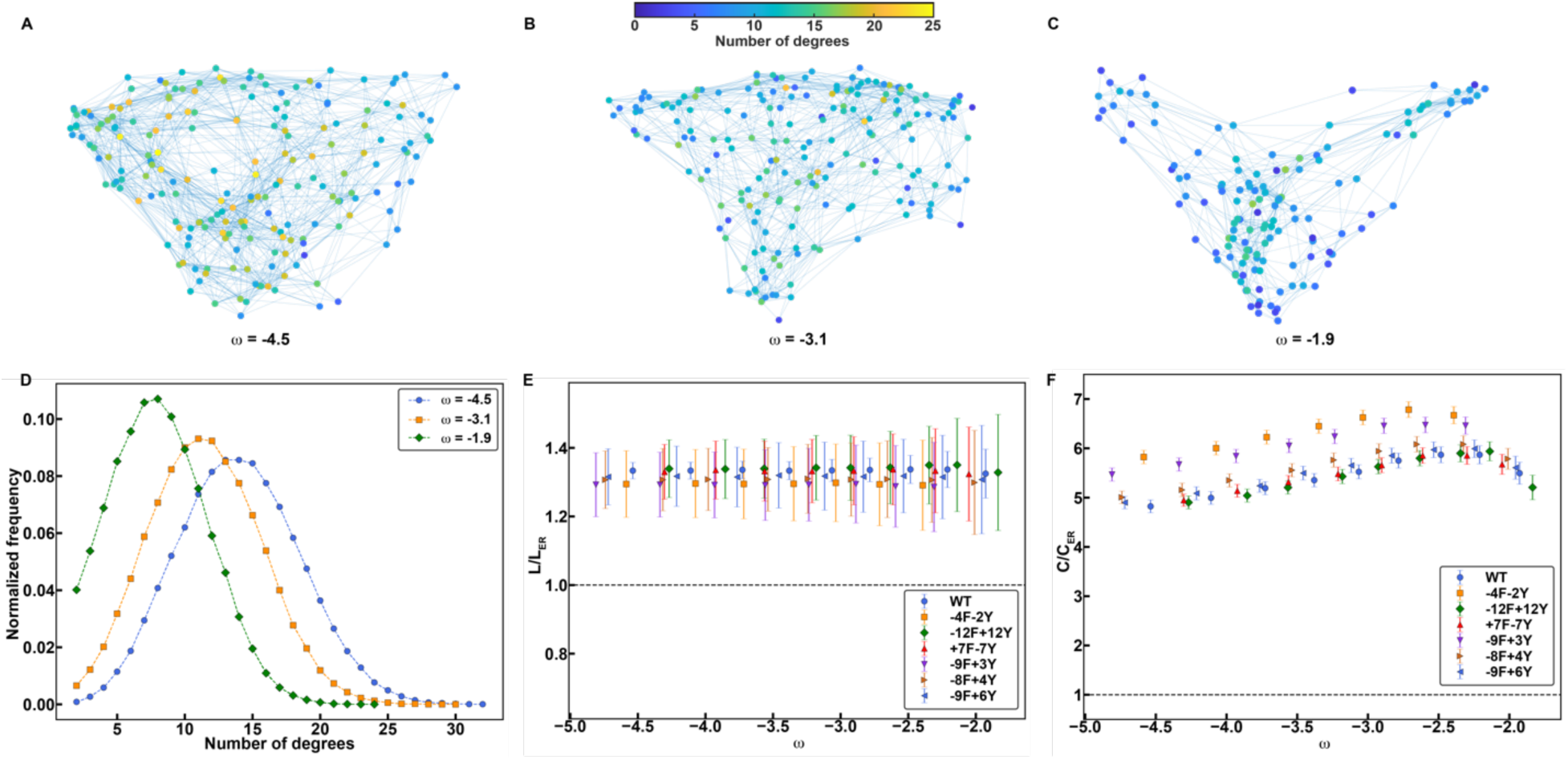
The interiors of condensates form small-world network structures. **(A-C)** Representative graphs for the largest connected cluster at the largest **(A)**, median **(B)**, and smallest **(C)** value of ω (as defined in Fig. 2). Results are shown here from analysis of simulations for the wild-type A1-LCD. The nodes represent individual molecules and are colored according to their degree (number of connections that they form). Two chains are connected by an undirected edge if any two stickers between them are within 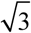 units on the cubic lattice. **(D)** The degree distribution for three distinct values of ω. **(E, F)**, the average path length, *L* **(E)**, and the average clustering coefficient, *C* **(F)**, for the wild-type A1-LCD and the six “aromatic variants”. The values shown here are normalized by the corresponding Erdős-Rényi values for random graphs. The dashed horizontal line represents the values that would be expected assuming an Erdős-Rényi model^40^. The error bars represent the standard deviation. As ω approaches zero, we note a downward shift of the relative clustering coefficient. This is indicative of increased randomness as the critical temperature is approached.

**Figure 4:**
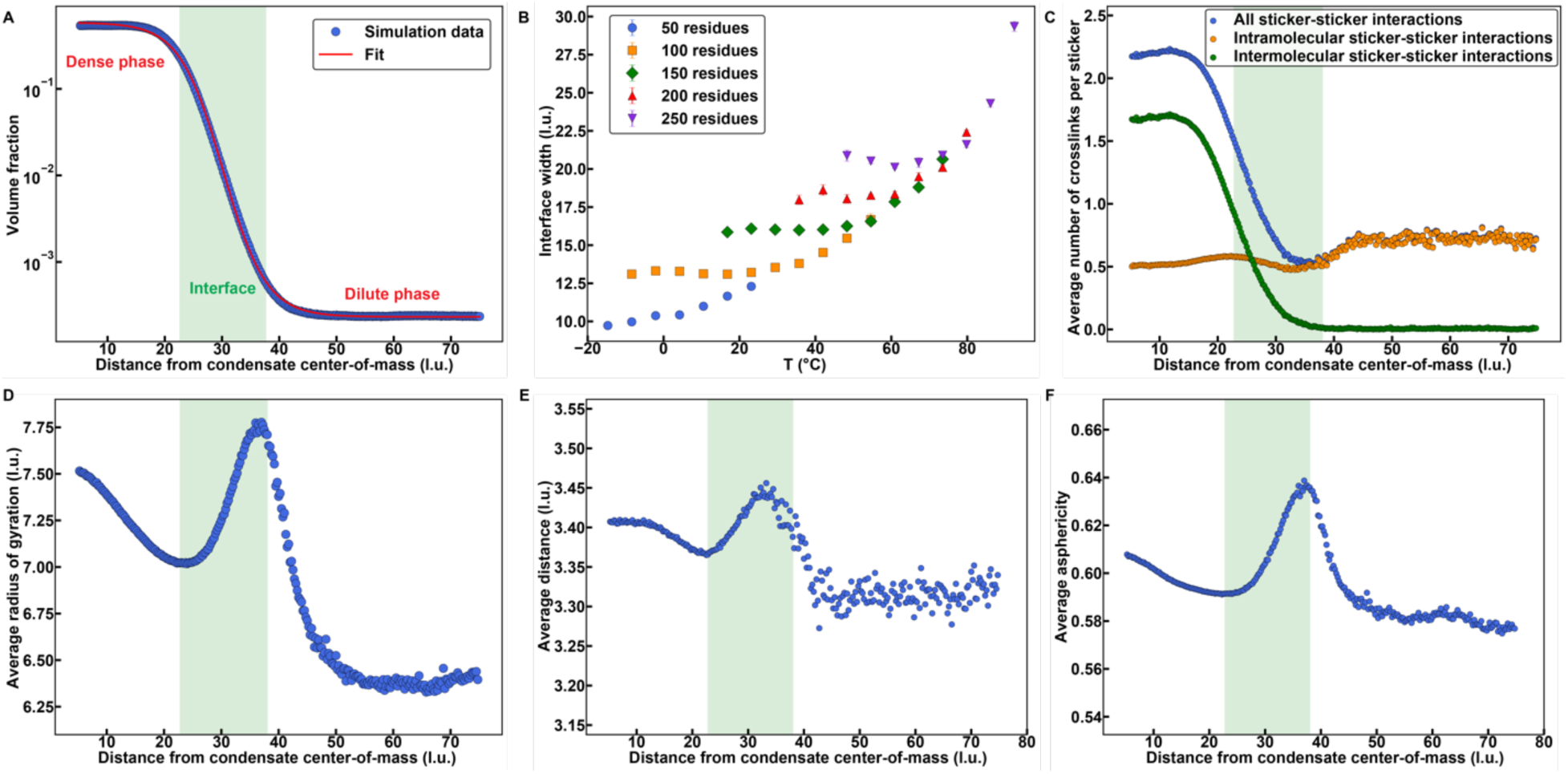
Interfaces of condensates have distinctive conformational characteristics. **(A)** A representative radial density plot of a simulation of the wild-type A1-LCD at ω=-3.7. The solid red curve corresponds to a logistic fit to the data (see Methods). **(B)** The width of the condensate interface versus temperature for simulations of homopolymers at different lengths. Error bars are standard errors about the mean across three replicates. **(C)** The average number of sticker-sticker crosslinks per sticker as a function of the distance from the condensate center-of-mass for wild-type A1-LCD at ω=-3.7. Depicted are the total number of crosslinks (blue), the number of intramolecular crosslinks (orange), and the number of intermolecular crosslinks (green). **(D)** The average *R*_*g*_ of a chain as a function of the distance from the condensate center-of-mass for the wild-type A1-LCD at ω=-3.7. **(E)** Average distance between residues on the same chain that are separated by exactly five residues plotted against the distance from the condensate center-of-mass of one of the residues for the wild-type A1-LCD at ω=-3.7. **(F)** Average asphericity of chains plotted against the distance from the condensate center-of-mass of the chain for the wild-type A1-LCD at ω=-3.7. Values of asphericity that are larger than 0.4 point to cigar-shaped conformations, at least on the local level ^52^. The distinction of chain dimensions across the dilute, dense, and interfacial regions disappears as the critical temperature is approached. The translucent green boxes in panels **(A), (C), (D), (E)**, and **(F)** represent the interfacial region as determined by the logistic fit. In all panels, l.u. is lattice units.

The degree distributions are unimodal, and the mean and mode shift toward smaller values as *T* increases i.e., as the magnitude of ω decreases (Fig. 3D). The degree distribution is broad and features nodes that have degrees that are well above the mean. This, as depicted in the topological structures shown in Figures 3A-C, suggests a small-world structure of percolated networks within condensates. To test this hypothesis, we computed two standard measures of graph topology, the relative path lengths, and relative clustering coefficients of condensate graphs, at different temperatures, by referencing these parameters to values obtained from Erdős-Rényi random graphs ^40^. The mean path length is defined as the average shortest path between all possible pairs of nodes on the graph. The clustering coefficient is a measure of the degree of clustering of the nodes on the graph. The mean path lengths of condensate graphs are larger than those of Erdős-Rényi graphs ^40^ (Fig. 3E), and the mean cluster coefficients are at least a factor of five larger for condensate graphs. These features highlight the non-random, inhomogeneous, small-world nature of condensate graphs wherein a few molecules make up hubs in the network, and the rest of the molecules are connected to these hubs via sticker-mediated physical crosslinks.

At first glance, the observed small-world structure is surprising given that all the molecules within the condensate are identical to one another. It appears that the combination of sticker-and-spacer architectures, the valence, strength of stickers (Fig. 5F), conformational heterogeneity which affects the interplay of intra-chain, inter-chain, and chain-solvent interactions (Extended Data Fig. 5), and the spatial location of a molecule with respect to the center of the condensate / interface are all factors that will contribute to the observed small-world structure. In each snapshot, some molecules become hubs that enable the formation of condensate-spanning networks.

**Fig. 5:**
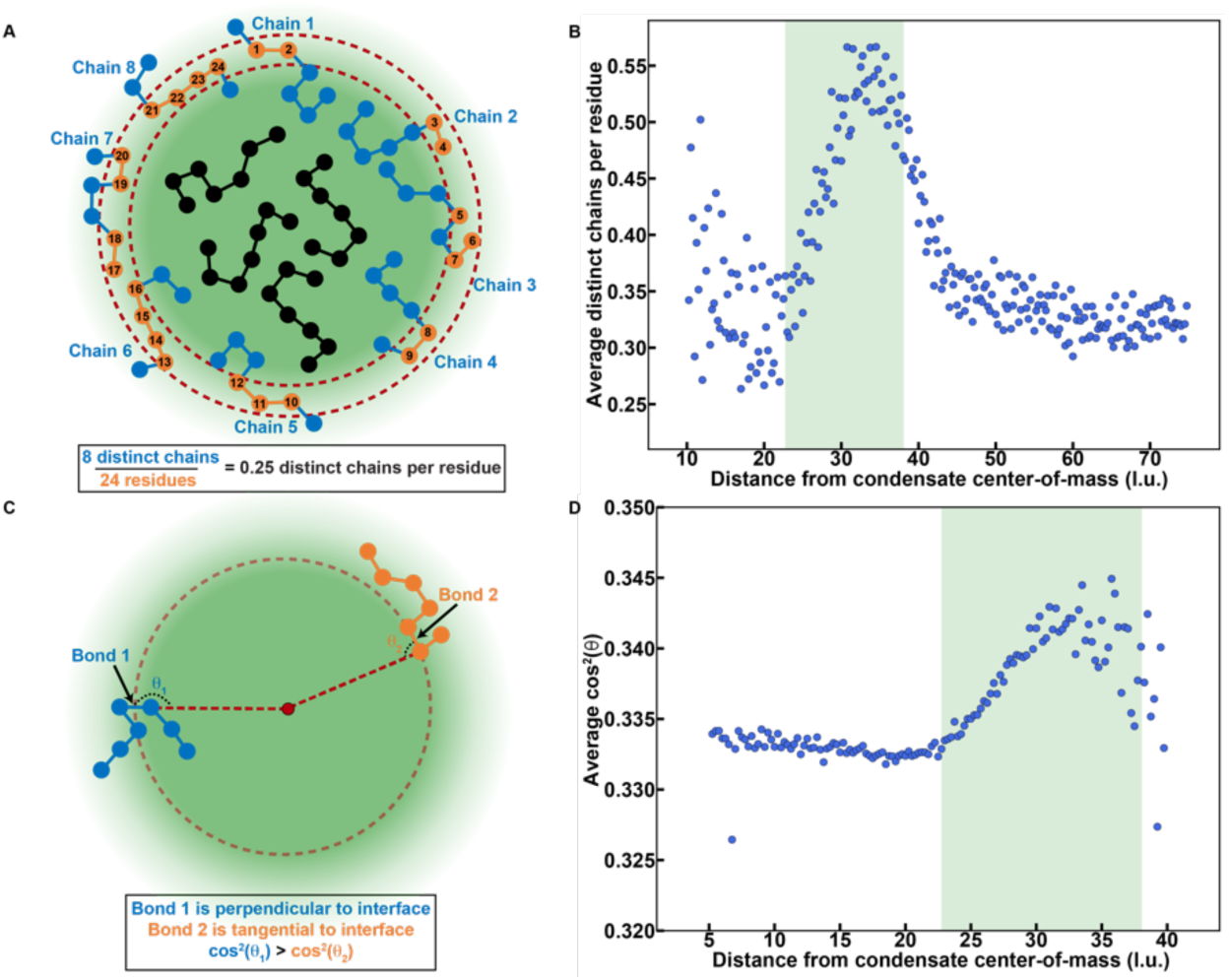
At the interface, molecules have non-random, perpendicular orientations. **(A)** A diagram depicting how distinct chains per residue is calculated. The region enclosed by the dashed red curves indicates the radial shell of interest. Any chains that contain beads within the radial shell are colored blue. Any beads that are within the radial shell are colored orange. All other chains are colored black. To calculate the distinct chains per residue, the number of blue chains is divided by the number of orange beads, in this case 8 and 24, giving a parameter value of 0.25. This parameter can vary between 0 and 1. Lower values suggest that chains are wrapped around a radial shell, whereas higher values suggests that chains are oriented perpendicular to a radial shell. **(B)** Average distinct chains per residue plotted against the distance from the condensate center-of-mass for the wild-type A1-LCD at ω=-3.7. **(C)** A diagram depicting how the parameter cos^2^θ is calculated. Here, θ is defined as the angle swept out by a covalent bond and a line segment (opaque dashed red line) between one of the bonded beads and the condensate center. Bond 1 (blue polymer) is perpendicular to the radial shell depicted by the translucent dashed red curve. Therefore, θ_1_ is close to 180° and cos^2^θ_1_ ≈ 1. Conversely, bond 2 (orange polymer) is tangential to the radial shell such that θ_2_ is close to 90°, and cos^2^θ_2_ ≈ 0. In general, lower values of cos^2^θ suggest that chains are wrapped around a radial shell, whereas higher values suggest that chains are oriented perpendicular to a radial shell. **(D)** Average cos^2^θ plotted against the distance from the condensate center-of-mass for the wild-type A1-LCD at ω=-3.7. Notice the increase in cos^2^θ values within the region corresponding to the interface. As the critical temperature is approached, the orientational differences across distinct regions vanish. Translucent green rectangles in **(B)** and **(D)** represent the interfacial region determined by the logistic fit in Fig. 4A. l.u. is lattice units.

The observed small-world network structures imply that even within condensates formed by molecules of a single type, the crosslinking density will be inhomogeneous. This can give rise to time-dependent changes of material properties, expected for viscoelastic materials, and physical aging ^41^, as has been observed for simple condensates such as those formed by PLCDs ^42, 43^and other low complexity domains ^44^. Additionally, the type of small-world network that is formed, as defined in terms of the degree, mean path length, and mean clustering coefficient, will be affected by solution conditions (temperature in our case), and the linear patterning of stickers ^41^. Recently, Shillcock et al.,^45^ used a specific implementation of graph-theoretical approaches to analyze their simulations of condensates formed off a lattice, using dissipative particle dynamics, for generic sticker-and-spacer models. They concluded that the network topologies within condensates have small-world architectures. Taken together, it appears that the small-world structure of condensates might be a feature of all linear sticker-and-spacer systems.

## Molecular features of condensate interfaces

In the two-phase regime, there exists an interface between coexisting dilute and dense phases. We analyzed radial density profiles to identify interfacial regions (Fig. 4A). Each radial density profile has two shoulders corresponding to coexisting regions of low and high densities. The density in the transition region changes monotonically between the two shoulders. This is the presumed interface between the coexisting dilute and dense phases. The interface will be defined by the wavelength of capillary fluctuations, the sizes of molecules at the interface, the surface density of molecules, and the orientations of molecules with respect to the interface ^46, 47^. Following precedents for describing liquid/vapor interfaces in van der Waals fluids and associative molecules ^48, 49, 50, 51^, we use a hyperbolic tangent function^49, 51^ to fit the computed radial density profile ϕ(*r*) at a given temperature. The function used is shown in Equation (2):

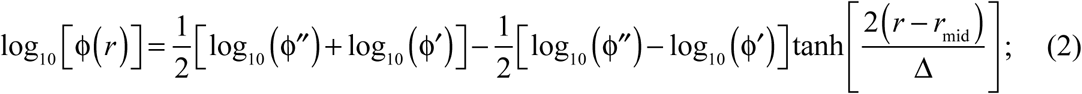

Here, ϕ′and ϕ″ are the densities in the dilute and dense phases, respectively; *r*_mid_ is the midpoint of the hyperbolic tangent function, and Δ is the inferred width of the interface. As shown in Fig. 4A, the computed radial density profile can be well described by the hyperbolic tangent function. We used this function to analyze how the width of the interface (Δ) scales with chain length (*N*) for homopolymers that were modeled using the parameters obtained to reproduce the measured and computed binodals of the wild-type A1-LCD (Extended Data Fig. 6). The width increases with temperature (Fig. 4B). Further, away from the critical temperature we observe a plateauing of Δ to a length-specific value Δ_p_, where Δ_p_ ∼ *N*^0.45^. This implies that the width of the interface increases with increasing molecular weight of the flexible polymer. Interestingly, above a length-specific temperature, as the temperature approaches *T*_*c*_, the width of the interface (Δ), which continues to increase, becomes independent of chain length.

Next, we analyzed the progression of inter-sticker contacts along the radial density profile (Fig. 4C). We observe a monotonic decrease in the average number of intermolecular, inter-sticker interactions along the radial coordinate *r* that progresses from the dense phase into the dilute phase (Fig. 4C). However, the average number of intramolecular, inter-sticker interactions changes non-monotonically. This value, which is low in the dense phase, decreases further through the interface, followed by an increase as *r* extends beyond the interface into the dilute phase (Fig. 4C). The conformational consequences of this non-monotonic change in intramolecular crosslinks per sticker are summarized in Figures 4D-F. As shown in Fig. 4D, the *R*_*g*_ values of individual molecules are largest within the interface and smallest within the dilute phase. The preference for expanded conformations is also manifest on local length scales as shown in Fig. 4E. Here, we demonstrate that sections of the chain that are up to five bonds long are generally more expanded at the interface when compared to the dense and dilute phases. The global expansion results from more prolate-shaped conformations^52^, as is shown by the evolution of the average asphericity^52^ along the radial coordinate (Fig. 4F). Overall, the results in Fig. 4 show that the width of the interface, even away from *T*_*c*_, is approximately three times larger than the average *R*_*g*_ of chains in the dilute phase. This implies that the width of the interface is at least as large as the mean end-to-end distance of a flexible PLCD. This observation is consistent with inferences reported in a recent study by Böddeker et al., ^53^ of condensates being defined by “fat” interfaces.

## Chains are oriented normally at the condensate interface

The increased global and local expansion we observe on average for molecules at the interface raises two possibilities for the orientations of molecules. First, they could be expanded because they adsorb and are oriented parallel to the interface. This arrangement would minimize the number of chains per unit area, ensuring that un-crosslinked stickers at the interface originate from a small number of distinct chains for a given condensate size. Alternatively, the chains could have a locally perpendicular orientation with respect to the interface. This arrangement would maximize the number of chains at the interface while minimizing the number of unsatisfied stickers per chain. We computed the average number of distinct chains per residue (Fig. 5A), resolved along the radial coordinate pointing from the center of the condensate. This value is maximized at the interface (Fig. 5B), implying that molecules do not adsorb, and are not oriented parallel to the interface. Instead, each chain section is oriented perpendicularly to the interface. To further test for this, we computed the projection angles of bond vectors with respect to the radius vector with origin at the center of the condensate that is being analyzed (Fig. 5C). Resolved along the radial coordinate, we note that the bond vectors prefer perpendicular orientations at the interface and random orientations within condensates (Fig. 5D).

## The dilute phase crosses over into the semidilute regime as *T* approaches *T*_*c*_

We find that on a semi-log scale, the dilute arm of the binodal shifts rightward with increasing temperature, whereas the dense arm shows little change (Extended Data Fig. 3). This implies that the width of the two-phase regime shrinks almost entirely because of an increase in the saturation concentration with temperature. Note that PLCDs have upper critical solution temperatures ^9^. In polymer solutions, there exists a special concentration that equals the concentration of chain units within the pervaded volume of a single chain ^54^. This is known as the *overlap concentration c\** - so named due to the high likelihood that chains will overlap with one another when the solution concentration exceeds *c*\* ^55^. In dilute solutions, *c* < *c*\*, whereas in semi-dilute solutions, *c*≈*c*\*. We used the mean end-to-end distance values in the single-chain limit ^56^ to compute temperature-dependent overlap volume fractions ϕ^*^(*T*) for the wild-type A1-LCD. For temperatures below 20°C, ϕ_sat_(*T*) < ϕ^*^(*T*) i.e., the left arm of the binodal is located to the left of the overlap line (Extended Data Fig. 8A). Accordingly, for *T* < 20°C, the dispersed phase that coexists with the dense phase is a true dilute phase. However, we observe a crossover above ∼20°C whereby ϕ_sat_(*T*) > ϕ^*^(*T*), which is caused directly by the increased density within the dilute phase (compare Extended Data Figs. 8B vs. 8C). Therefore, the dispersed phase that coexists with the condensate is semi-dilute for temperatures above ∼20°C. These distinctions are relevant because properties of polymer solutions in dilute solutions are governed exclusively by the interplay of intramolecular and chain-solvent interactions. Conversely, the physical properties of semi-dilute solutions are governed by the interplay of density fluctuations and conformational fluctuations, which impacts intramolecular, intermolecular, and chain-solvent interactions ^54, 55^. Further, the dynamics of individual chains and overall rheological properties of dilute and semi-dilute solutions will also be considerably different from one another, with increasing viscoelasticity characterizing semi-dilute solutions.

## Overall implications of our findings

We have built upon recent experimental characterizations of phase behaviors of the A1-LCD system and designed variants thereof to develop a lattice-based, single bead per residue model that accurately captures the measured phase diagrams. It is noteworthy that the data of Bremer et al.,^9^ and those on unrelated low complexity domains have also been qualitatively and quantitatively reproduced by other, off-lattice coarse-grained models ^57, 58^. A specific approach used to compare computations and experiments is through the comparison of computed vs. experimentally derived critical temperatures ^57^. However, estimates of *T*_*c*_ are inaccessible from direct measurements. They are instead inferred by fitting binodals extracted from a preferred mean-field theory, cf., Extended Data Fig. 8A. Then, a specific functional form for the width of the two-phase regime ^33, 57, 59^ is fit to the entire binodal. However, based on the Ginzburg criterion, the functional form that is routinely used ^33, 57, 59^ is only valid in the vicinity of the critical temperature ^60^. We pursue a different approach to compare computed and experimentally derived phase diagrams. Specifically, we quantify the ERMSL between computed and measured low concentration arms of binodals. We focused on the low concentration arms because they change the most with temperature and have smaller error bars in measurements when compared to concentrations corresponding to the high concentration arms of binodals. Overall, the ERMSL values indicate that maximal deviations are a factor of 2-2.5 across concentrations that vary by at least three orders of magnitude. Encouraged by the accuracy of our simulations across 31 different variants of the A1-LCD system, we used the computed ensembles within, outside, and at the interface of condensates to investigate molecular and mesoscale structures.

Our findings regarding the degree of crosslinking and extent of chain expansion in the three regions, *viz*., dilute phase (I), condensate interface (II), and condensate interior (III), are summarized in Fig. 6. Our results suggest that interfaces between condensates and the coexisting dilute phases should be thought of as being “fat” ^53^ rather than “thin”. This feature of the interface is realized by the ability of disordered proteins to be relatively more expanded, both locally and globally, when compared to the dilute and dense phases. It is known that the interfacial tension decreases as the inverse square of the size of the molecule ^46^. Accordingly, the low interfacial tensions that have been measured to date ^61 42, 62, 63^ appear to originate from chains being most expanded as they traverse the interface. Importantly, the interface features a high number of unsatisfied stickers, achieved due to the high number of chains that project perpendicularly to the interface.

**Fig. 6:**
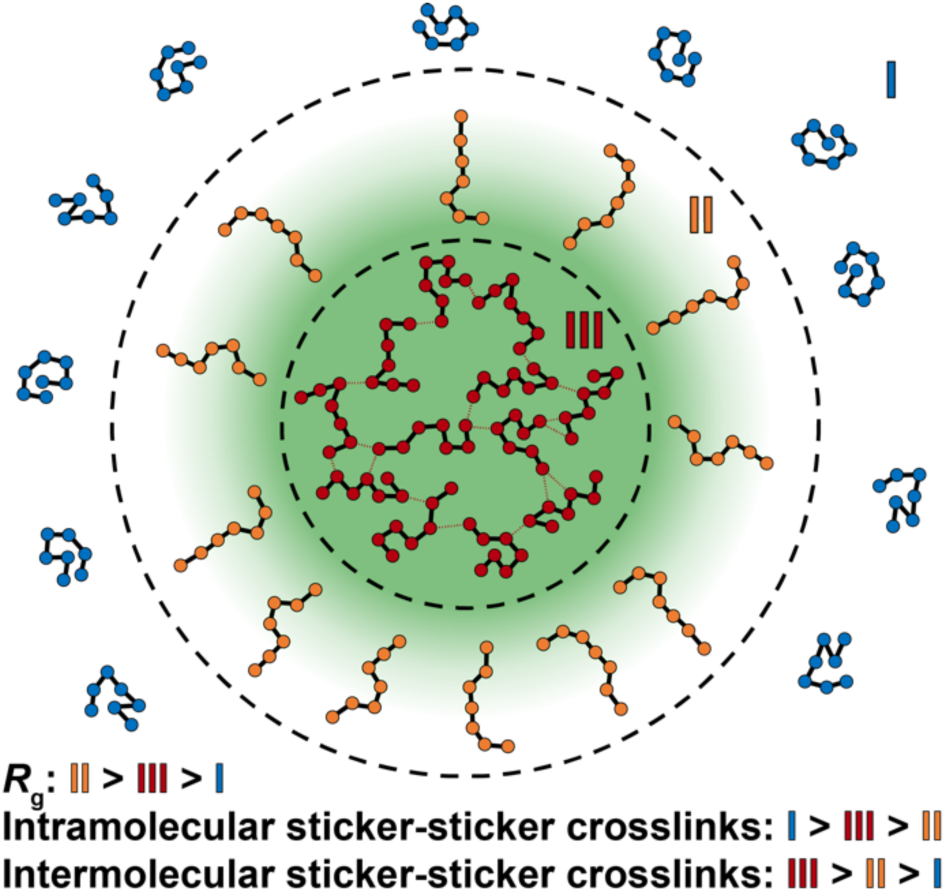
Molecular properties of interfaces are distinct from dilute and dense phases. Diagram summarizing our findings concerning condensate organization. Region I is the dilute phase, Region II is the condensate interface, and Region III is the interior of a condensate. Region I is characterized by relatively compact chains that form few intermolecular contacts. Region II is characterized by relatively expanded chains that are oriented perpendicular to the interface and form the fewest number of total sticker crosslinks. Region III is characterized by chains that are less compact than those in Region I and less expanded than those in Region II. These chains form numerous intermolecular sticker crosslinks, giving rise to a small-world percolated network.

Our observations regarding interfaces have two major implications. First, the presence of a high number of unsatisfied stickers, typically defined by the presence of functional groups, suggests that interfaces might be prime locations for enhancing the efficiencies of biochemical reactions that are influenced by condensate formation. This speculation, based on the features we have documented for interfaces, is consistent with numerous observations from the microdroplet literature ^64^. Second, it is conceivable that interfaces catalyze amyloid fibril formation, through secondary nucleation ^65, 66^. This proposal is based on the preference for the high likelihood of accessing locally extended, β-strand-like conformations for molecules such as A1-LCD or mutations of such systems ^43^. Our proposal appears to be supported by the recent results of Linsenmeier et al., ^67^ who report that amyloid formation is nucleated at condensate interfaces.

Overall, our findings suggest that even the simplest condensates, formed via effective homotypic interactions among PLCDs with sticker-and-spacer architectures, have complex internal structures and interfacial characteristics. The features we have identified are likely to be germane to recent discoveries that condensates are in fact viscoelastic network fluids ^68, 69^. We find that condensates formed by PLCDs have small-world network structures. This implies that the molecules are organized into inhomogeneous networks within condensates defined by regions of high vs. low crosslinking densities. The clustering of stickers within condensates, achieved via strong and specific inter-sticker interactions, can be separated from the contributions of spacers that directly impact the solvation preferences, thereby modulating the locations of the dilute arms of binodals. The extension of our findings to multicomponent, multiphasic systems ^70^ will be of considerable ongoing and future interest.

## Methods

### Monte Carlo simulations using LaSSI

Simulations were performed using LaSSI, a lattice-based Monte Carlo simulation engine ^20^. Monte Carlo moves are accepted or rejected based on the Metropolis-Hastings criterion such that the probability of accepting a move is the min[1,exp(-Δ*E*/*k*_*B*_*T*)] where Δ*E* is the change in total system energy of the attempted move and *k*_*B*_*T* is the thermal energy. Total system energies were calculated using a nearest neighbor model whereby any two beads that are within one lattice unit of each other along all three coordinate axes contribute to the total energy of the system. In our case, we define all pairwise interaction energies as absolute energies (Fig. 5B) that are scaled by the simulation temperature during the Metropolis-Hastings step.

### Calculation of phase diagrams and interfacial features

Multi-chain LaSSI simulations at various temperatures were performed to calculate coexistence curves. For each variant, 200 chains, each with 137 beads, were placed in a cubic lattice with side-length 120. The starting volume fraction was about 0.016. After the systems phase separated, the radial distribution of beads in the system was calculated as shown in Equation (2) of the main text. All multi-chain simulations were performed in triplicate, and the calculated volume fractions were near identical across replicates. At low temperatures, the calculated dilute phase volume fractions tend to have more variability due to the decreased likelihood that chains are in the dilute phase. To convert from simulation temperature and volume fraction to experimental temperature and volume fraction, fixed scaling factors of 5.6 and 0.6 were used ^8, 9^, respectively. These scaling factors were used for all variants, and they were chosen by comparing the simulation and experimental phase diagrams of the aromatic variants (Extended Data Fig. 3A-B). In general, scaling factors are required to provide phase diagrams that match experimental results. To convert from volume fraction to mass concentration, we assumed that a volume fraction of 1.0 corresponds to a mass concentration of 1310 mg/ml ^56^.

### Gaussian process Bayesian optimization (GPBO) for parameterization and verification of sticker-and-spacer model

Single-chain LaSSI simulations at a single temperature (*k*_*B*_*T* = 50) were performed to parameterize the initial stickers-and-spacers model. Values of the apparent scaling exponent (ν_app_) derived from experimental SEC-SAXS data of the variants in Extended Data Fig. 3A-E of Bremer et al.,^9^ were used as the target values. Values of ν_app_ were calculated as described by Meng et al., ^71^ using ten independent simulations per construct. A Gaussian process Bayesian optimization ^72^ was implemented. This process was iterated over pairwise interaction energies to minimize the sum of the square residuals of computed and experimentally derived values for ν_app_ (Fig. 1B). If a given parameter remained close to the upper or lower bound through each iteration, then the parameter bounds used in the optimization were manually changed over the course of the optimization. Over 500 iterations were performed. The final parameters as well as the upper and lower bounds and values are shown in Table 1. Here, “Aro” represents either a Tyr or Phe residue and “X” represents any amino acid residue that is not explicitly one of the residues listed in the main text. We note that for every parameter, the difference between the final value and either of the bounds is at least 20% of the absolute final value, suggesting the bounds are adequately distanced.

**Table 1:** Initial parameterization of sticker-and-spacer model using GPBO.

| Pairwise Interaction | Lower Bound | Upper Bound | Final Value |
| --- | --- | --- | --- |
| Tyr-Tyr | -32 | -15 | -19.3 |
| Tyr-Phe | -30 | -12 | -15.2 |
| Phe-Phe | -25 | -5 | -13.4 |
| Arg-Aro | -25 | -8 | -11.0 |
| Lys-X | -3 | 3 | -0.389 |
| X-X | -4 | -2 | -2.95 |

### A mean-field model to account for protein charge

We have previously shown that small changes to the net charge per residue (NCPR) have little effect on single chain dimensions but can alter dilute phase concentrations by orders of magnitude ^9^. To account for this, we incorporate a mean-field NCPR-based term into our multi-chain simulations. This term strengthens or weakens the interactions of a system, by decreasing or increasing the effective temperature, respectively, depending on the NCPR of the variant. In this way, we implicitly account for ionizable residues without explicitly modeling them as unique, highly solvated spacers. The mean-field term is incorporated as follows:

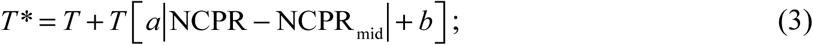

Here, *T*\* the new effective temperature, *T* is the original temperature, NCPR is the net charge per residue of the variant of interest, and NCPR_mid_, as well as *a* and *b* are constants. The mean-field term can be visualized as an absolute-value function whose minimum vertex is at (NCPR_mid_, *b*) and whose slope is -*a* to the left of the minimum and +*a* to the right of the minimum (see Extended Data Fig. 2). Based on prior work, NCPR_mid_ = 0.0287, *a* = 1.54, and *b* = -0.045. This ensures that *T*\* = *T* for the wild-type A1-LCD.

### Parameterization of spacer interactions

Although we did not have SAXS data for the variants designated as “spacer variants” by Bremer et al.^9^, we were able to parameterize interactions between spacers and other beads by manually titrating five new interaction parameters to match the computational phase diagrams to the experimental phase diagrams shown in Extended Data Fig. 3F-I. Parameters for the final model are shown in Fig. 1A and Table 2. In cases where a pairwise interaction is ambiguous (for example, the interaction between Thr and Ser), the weaker interaction is used (in this case, -2.35).

**Table 2:**
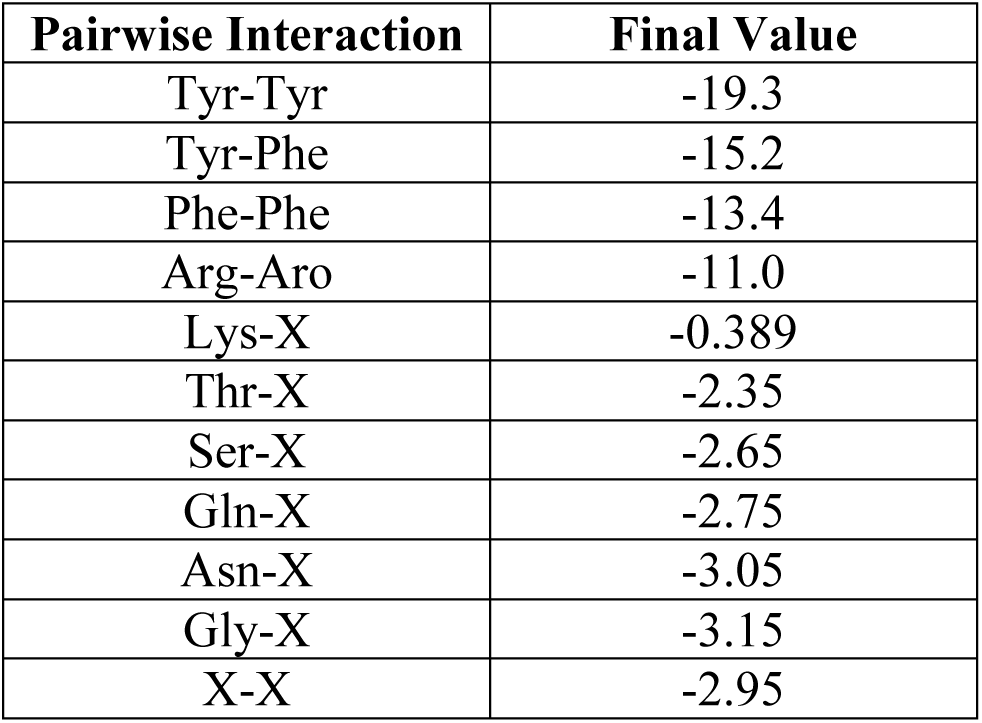
Final parameterization of sticker-and-spacer model.

### Error analysis of computed phase diagrams

We evaluate the error of our LaSSI-derived phase diagrams from the experimentally derived phase diagrams using a multi-step process: First, we perform a linear regression of temperature vs. log(c_sat_) for the LaSSI-derived dilute arms. (2) Second, for each experimentally-derived data point along the dilute arm, calculate *x* = (*c*_sat,sim_ / *c*_sat,exp_) where *c*_sat,sim_ and *c*_sat,exp_ are the computed and experimentally derived saturation concentrations at the same temperature. Third, we calculate the exponential root mean square log (ERMSL) of *x* as 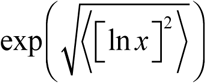. The ERMSL is a positive value greater than or equal to 1 and can be interpreted as a measure of the error between the computed and experimentally derived phase diagrams, specifically the dilute arms. For example, an ERMSL value of 10 indicates that, on average, the two *c*_sat_ values differ by about a factor of 10, or one order of magnitude. Alternatively, an ERMSL value of 1 indicates that there is no error between the dilute arms and that they should overlay perfectly. We report the ERMSL of every variant in Fig. 1D. This includes the variants in Extended Data Fig. 3J, which were not used in the parameterization of the sticker-and-spacer model. The ERMSL is akin to the root mean square log error (RMSLE) often used in machine learning, except for two important distinctions: (1) When calculating the RMSLE, 1 is added to both the numerator and the denominator of the argument. In our case, we do not need to include this bias since *c*_sat_ values are always greater than zero. (2) Unlike with the RMSLE, we take the exponential of our final value to bring our error back to an interpretable scale. This exponential operator is a reciprocal function of the inner logarithmic operator, in the same way that the outer square root operator is a reciprocal function of the inner square operator.

### Analysis of conformational properties in dense and dilute phases

In Fig. 2, Fig. 3, Extended Data Fig. 4, and Extended Data Fig. 5, we report conformational characteristics of PLCD molecules in dense and dilute phases. To perform these analyses, we first ensure that our simulation shows stable phase separation into a single, distinct dense phase. We then determine whether a chain belongs to the dense or dilute phase based on whether it is within interacting range of the largest cluster of chains. If so, we group this chain into the dense phase. Otherwise, we group this chain into the dilute phase.

### Swelling ratio

In Fig. 2 and Extended Data Fig. 4, we introduced the swelling ratio α, which for a given temperature or ω-value, we define as:

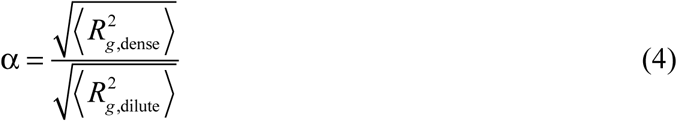

As noted in the main text, we defined the width of the two-phase regime in terms of the parameter ω. Rather than directly calculating the difference between the dense and dilute phase concentrations, which is heavily biased by the dense phase concentration, we calculate the difference between the concentrations on a log scale. This accounts for the fact that the dense and dilute phase concentrations differ by orders of magnitude. The swelling ratio quantifies the degree of expansion of chains in the dense phase relative to the dilute phase. In Fig. 2C, we fit the following exponential decay model to data for the swelling ratio:

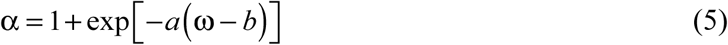

Here, *a* is the fitted parameter that controls the rate of decay, and *b* shifts the curve to the left or the right. The parameters for the master curve shown in Fig. 2C are *a* = 0.34, and *b* = -9.4, respectively.

### Ternary plots to analyze the interplay of intra-chain, inter-chain, and chain-solvent interactions in the dense phase

Extended Data Fig. 5 shows a ternary plot constructed in the following way: (1) For every chain in the dense phase, we calculate the *R*_*g*_ values of individual molecules. (2) For each bead in that chain, we count the number of neighbors within 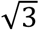 lattice units that are empty (i.e., solvated), contain beads that belong to the same chain, or contain beads that belong to other chains. (3) Sum each count for every bead in the chain and divide by the sum of the three counts. This yields the three fractions, *f*_sol_, *f*_intra_, and *f*_inter,_ respectively. (4) For each chain, we then determine which bin it belongs to on the ternary plot (based on the fractions) and average *R*_*g*_ calculated for all chains in that bin to determine the color of the bin of interest.

### Overlap concentration calculation

In Extended Data Fig. 8A, we calculate the overlap concentration of simulated constructs in a using the method of Wei et al., ^56^. Specifically, we use the following equation:

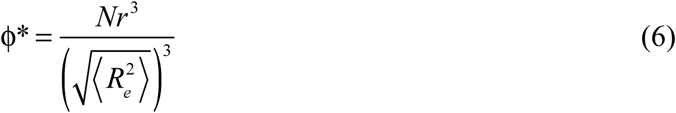

Here, ϕ* is the overlap concentration, *N* is the number of monomers in a chain, *r* is the radius of individual residues and 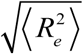 is the root mean square end-to-end distance of a chain. In our case, *N* is 137 and *r* is set to 0.5 lattice units. We apply this equation to chains in the dilute phase to minimize the effects of intermolecular interactions on the calculation of the overlap concentration dictated purely by conformational fluctuations.

### Parameterization of a model for an equivalent homopolymer

In Extended Data Fig. 6, we introduce a homopolymer equivalent for the wild-type A1-LCD. The contact energies for this model were parameterized by choosing a single pairwise interaction energy such that the phase diagrams of the homopolymer and wild-type A1-LCD overlay on one another. A pairwise interaction energy of -3.3 accomplishes this task.

### Graph theoretical analyses

In Fig. 3, we report on the small-world network structure formed by condensates. For this, we analyze the undirected, unweighted graphs formed by each condensate over the equilibrated portion of the trajectory ^39^. For each snapshot, we associate the condensate with the largest connected number of chains. Two chains are considered connected if at least one pair of stickers between them are adjacent, defined by being within 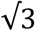 lattice units on the cubic lattice. For each condensate, we calculate the average path length and the average clustering coefficient to verify the small-world characteristics of the graph. The empirical average path length, representing the average number of steps along the shortest paths for all possible pairs of nodes (*v*_*i*_, *v*_*j*_), is calculated according to:

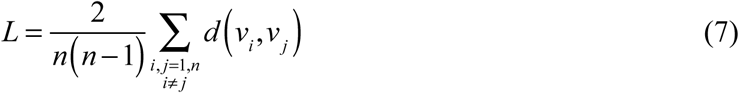

Here, *n* is the number of nodes. We compare this value to the average path length assuming Erdős-Rényi statistics ^73^, which we calculate as 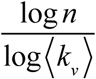 for each condensate, where ⟨*k*_*v*_⟩ is the average degree. Finite-size effects were not accounted for because ⟨*k*_*v*_⟩≪ *n*. We calculate the global clustering coefficient following the work of Watts and Strogatz ^74^ by averaging over the local clustering coefficients for all nodes. For an undirected graph, the local clustering coefficient is given by:

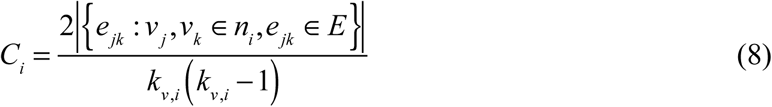

This calculation applies to nodes *v*_*j*_ and *v*_*k*_ that are in the neighborhood of node *n*_*i*_, with *e*_*jk*_ edges, in the set *E* of edges. The Erdős-Rényi value is calculated for each snapshot as 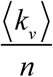.

### Analysis of internal organization of condensates

In Extended Data Fig. 7A-C, we report the likelihood that a sticker within a condensate is a neighbor of another sticker vs. a spacer, normalized by the same likelihood for the homopolymer. We calculate this parameter in the following way: (1) For a given condensate, we go through all the beads in each of the chains. (2) If a bead is a sticker (Tyr or Phe), we tally the number of its neighbors that are within 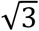 lattice units that happen to be stickers (*n*_st_) and the number of neighbors that are spacers (*n*_sp_). (3) We sum over all values to calculate *p*_a,seq_ (see main text). (4) We repeat steps 1-3 for the homopolymer condensate to calculate *p*_a,ref_. For this calculation, we assume that the homopolymer has the same sticker-spacer architecture as the wild-type A1-LCD. (5) The ratio of association *g*_a_ is then computed as shown in the main text. We note that Panel A and Panel C use wild-type A1-LCD (WT) as the background for the homopolymer, whereas Panel B uses WT+NLS, which includes one extra Tyr residue compared to WT.

### Weibull fits of sticker cluster probability distributions

In Extended Data Fig. 7D-F, we analyze the probability distributions for realizing clusters with stickers that form via inter-sticker crosslinks. For each equilibrated snapshot, we calculate the relative frequency that stickers form a cluster of a particular size, where the size is determined by the total number of stickers in the cluster. We then multiply each frequency by the cluster size to obtain the probability for a sticker to be in each cluster. A least-squares analysis, weighted by the inverse of the variance of repeated measurements, was performed on the linearized form of Equation 1. The analysis was restricted to the linear region of the plot. Outliers, where the distribution was exponentially bounded or where there were limited statistics at large *s* were treated as being a point that is greater than 3 scaled median absolute deviations from the median and hence removed from the analysis. The data treatment was insensitive to different outlier criteria. The values for the Weibull parameters were extracted directly from the fits to the linearized form of the cumulative distribution function given by Equation 1.

### Analysis of radial features to determine radial bins

Fig. 4, Fig. 5, and Extended Data Fig. 8 contain analyses of radial features of simulations. For each analysis, we use radial shells with thickness 1/4 of a lattice unit for the purpose of binning values together. In cases where we need a prior radial distribution, namely for determining the volume fraction in Fig. 4A and Extended Data Fig. 8B-C, we use the exact prior for a cubic lattice with side-length 120. When calculating the radial bins for the chains in Fig. 4D and Fig. 4F, rather than using the center-of-mass of a chain and counting each chain one time, we independently count each bead in the chain using the radial bin of the bead and the radius of gyration (Fig. 4D) or asphericity (Fig. 4F). This accounts for the fact that a single chain can span multiple bins by weighting each bin based on how many of a chain’s beads belong to it. Alternatively, in Fig. 4E, we only use the first bead in determining the radial bin.

### Average number of crosslinks per sticker

In Fig. 4C, we calculate the average number of crosslinks per sticker. To do so, we go through every sticker, defined as a Tyr or Phe residue, in the system and count how many of its neighbors within 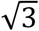 lattice units are also stickers. Each neighboring sticker represents one inter-sticker crosslink. We also determine whether each of these crosslinks is an intra- or intermolecular crosslink.

### Average distinct chains per residue

In Fig. 5A-B we describe and report a parameter which we term “average distinct chains per residue.” To calculate this parameter, we do the following: (1) for each radial shell, we count the number of distinct chains with beads that are contained in this shell. (2) Designate the number of distinct chains *n*_*c*_ and the total number of beads in the shell *n*_*b*_. (3) Calculate the final parameter as *n*_*c*_ / *n*_*b*_. This parameter is necessarily between 0 and 1. Lower values suggest that the beads in the given radial shell belong to a few distinct chains, whereas higher values suggest that the beads belong to many distinct chains.

### Orientational analysis

In Fig. 5C-D we describe and report a parameter that describes bond orientation relative to the condensate center-of-mass. To calculate this, we do the following: (1) for each covalent bond in the system, we consider a line segment drawn from one of the bonded beads (*b*) to the condensate center-of-mass. (2) We label the angle swept out by the covalent bond and the line segment in step 1 as *θ*. (3) Calculate cos^2^ *θ* and the radial bin to which *b* belongs. (4) For each radial bin, average the associated cos^2^ *θ* values. This parameter is necessarily between 0 and 1.

## Data availability

All simulation results and the details for reproducing the analyses are available via the Github repository of the Pappu lab.

## Code availability

All code used in this work is available via the Github repository of the Pappu lab.

## Acknowledgments

This work was supported by the US National Institutes of Health through grant R01NS121114 (T.M. and R.V.P), the St. Jude Children’s research collaborative on the biophysics and biology of RNP granules (T.M., and R.V.P), and grant FA9550-20-1-0241 from the Air Force Office of Scientific Research (R.V.P). We are grateful to Paolo Arosio, Furqan Dar, Matthew King, and Kiersten Ruff, for helpful discussions. We also acknowledge intellectual feedback from members of the Multidisciplinary University Research Initiative (MURI), Fundamental Design Principles for Engineering Orthogonal Liquid-Liquid Phase Separations in Living Cells that involves the labs of Jose Avalos, Cliff Brangwynne, Ashutosh Chilkoti, Amy Gladfelter, Rohit Pappu, and Lingchong You.

## Author contributions

M.F., and R.V.P. designed the simulations. M.F., S.R.C., and R.V.P., designed the analyses. W.M.B., A.B., and T.M., provided the experimental data on which the LaSSI model was parameterized. M.F., and S.R.C., analyzed the simulation results. M.F., S.R.C., and R.V.P., wrote the manuscript. All authors contributed to critical reading and editing of the manuscript.

## Extended Data Figures

**Extended Data Figure 1:**
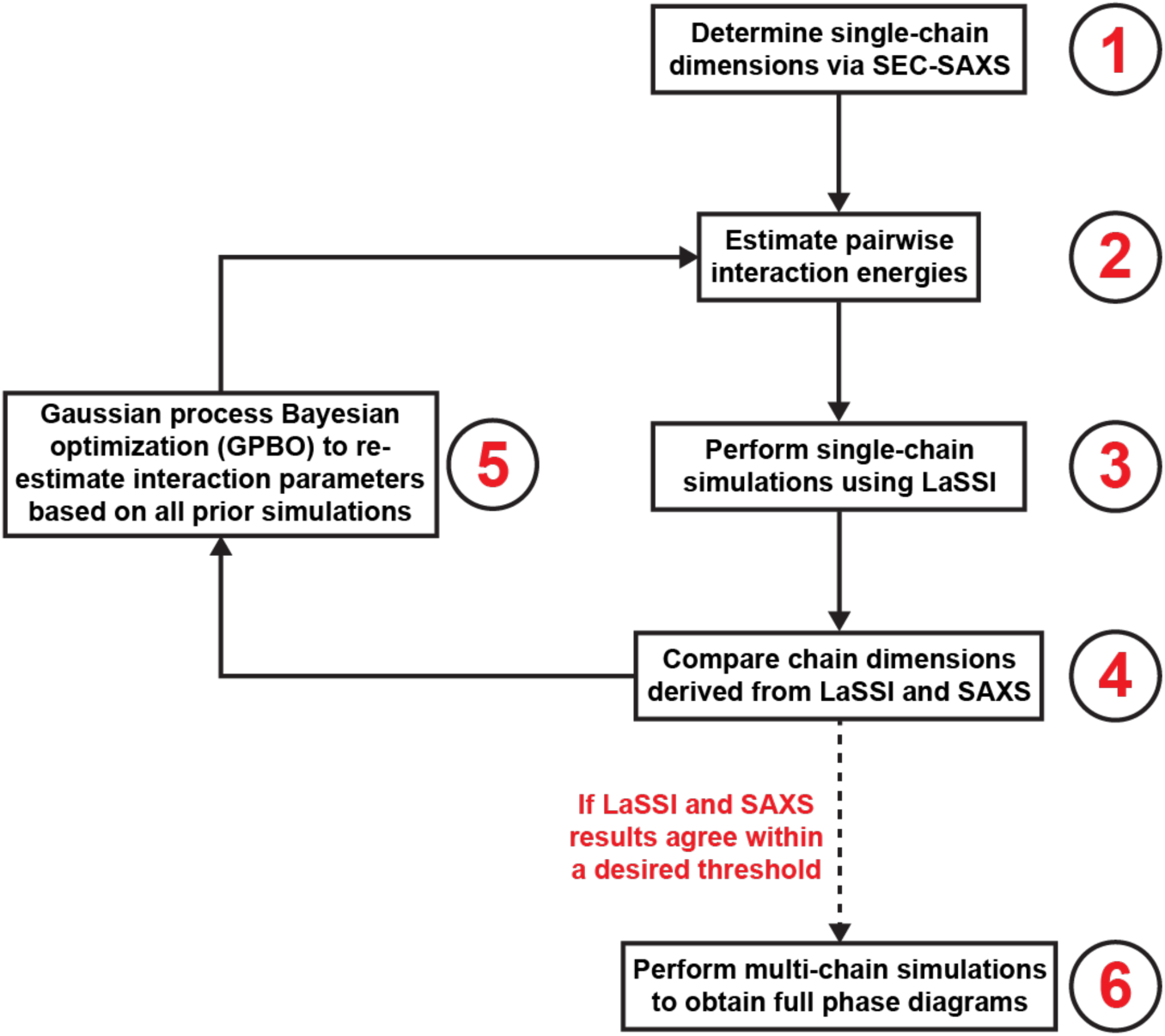
A flowchart describing the Gaussian process Bayesian optimization used to parameterize the computational model.

**Extended Data Figure 2:**
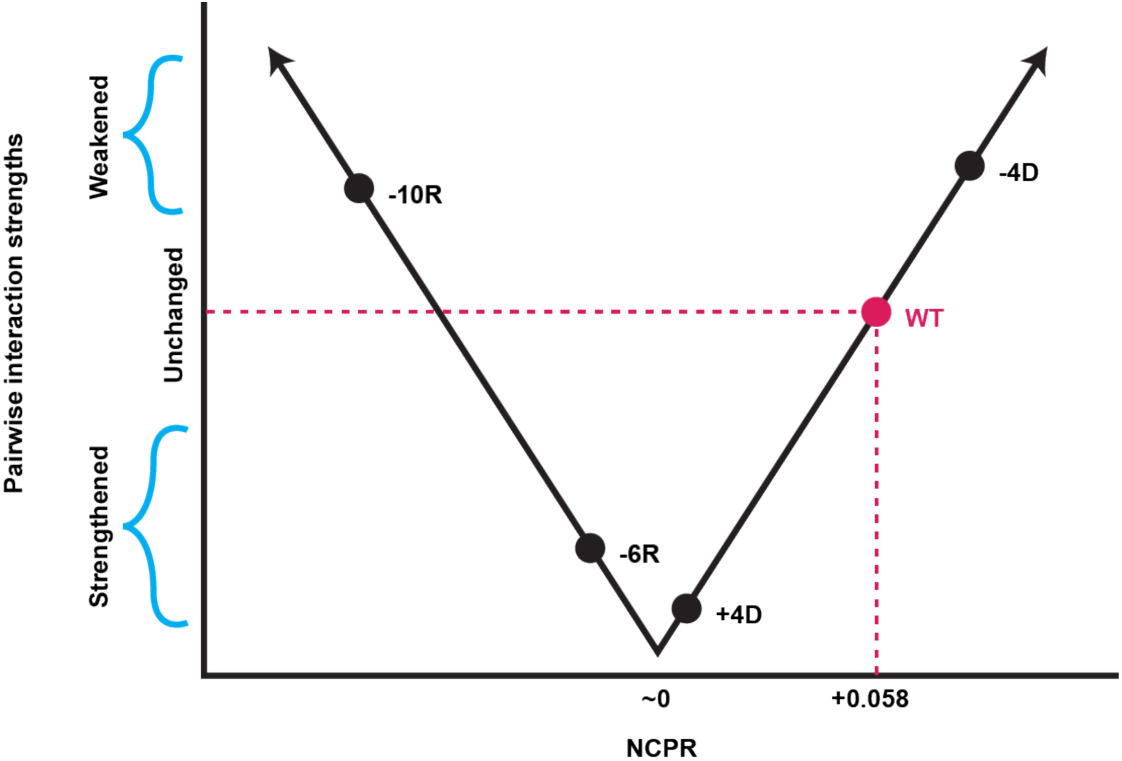
A diagram depicting how charge is incorporated in multi-chain simulations. In accordance with previous findings, a high magnitude of net charge per residue (NCPR) will cause a protein to be more soluble and phase separate at a higher concentration. In contrast, a low magnitude of NCPR will cause a protein to be less soluble and phase separate at a lower concentration. We take this into account by comparing the NCPR of a given variant with that of wild-type A1-LCD. As shown in this plot, the NCPR of wild-type A1-LCD is 0.058. If a variant has a significantly higher |NCPR| than 0.058, we weaken the pairwise interaction strengths among protein molecules. Similarly, if a protein has a lower |NCPR|, we strengthen the pairwise interaction strengths. Depicted are 4 other A1-LCD variants: -10R and -4D, whose |NCPR| values are higher than that of the wild-type, resulting in weakened pairwise interactions, as well as -6R and +4D, whose |NCPR| values are lower than that of the wild-type, resulting in strengthened pairwise interactions. See Methods for more details on this process.

**Extended Data Figure 3:**
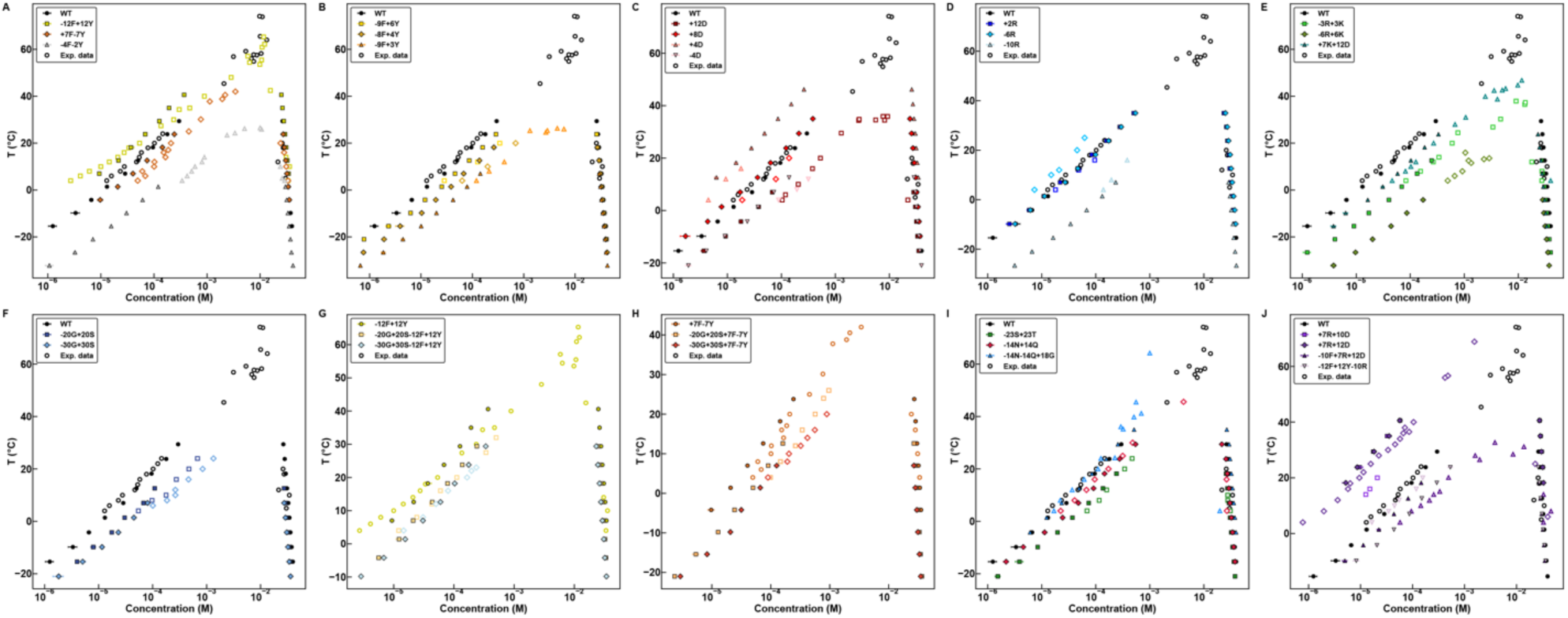
Calculated phase diagrams (solid markers) of all A1-LCD variants used in this study plotted alongside experimental phase diagrams (open markers). Temperature and concentration are converted from simulation units to experimental units by the same scaling factors for each variant. Variants depicted in the final panel (+7R +10D, +7R +12D, -10F +7R +12D, and -12F +12Y -10R) were not used to parameterize the computational model. Error bars indicate standard errors from the mean across 3 replicates.

**Extended Data Fig. 4:**
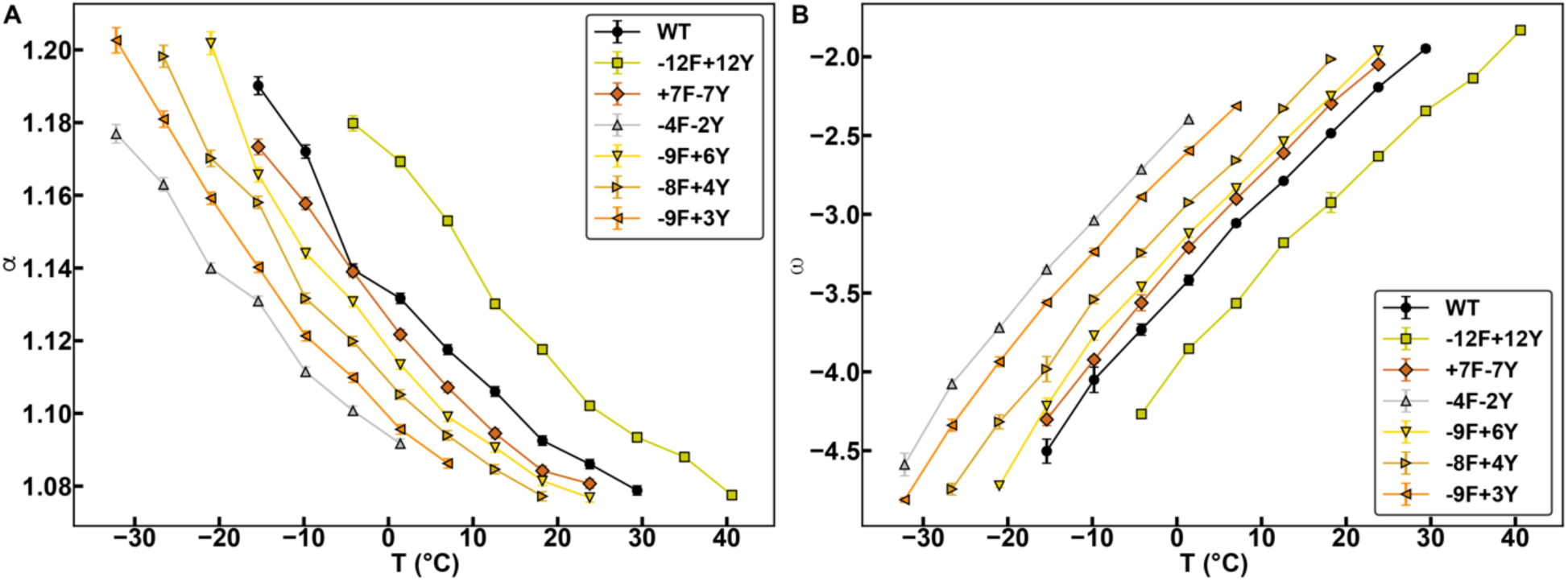
Swelling ratios and widths of two-phase regimes for different variants. **(A)** Average swelling ratio, α, plotted against temperature for aromatic variants of A1-LCD. **(B)** Average width of the two-phase regime, ω, plotted against temperature for aromatic variants of A1-LCD. Error bars indicate standard errors from the mean across 3 replicates.

**Extended Data Figure 5:**
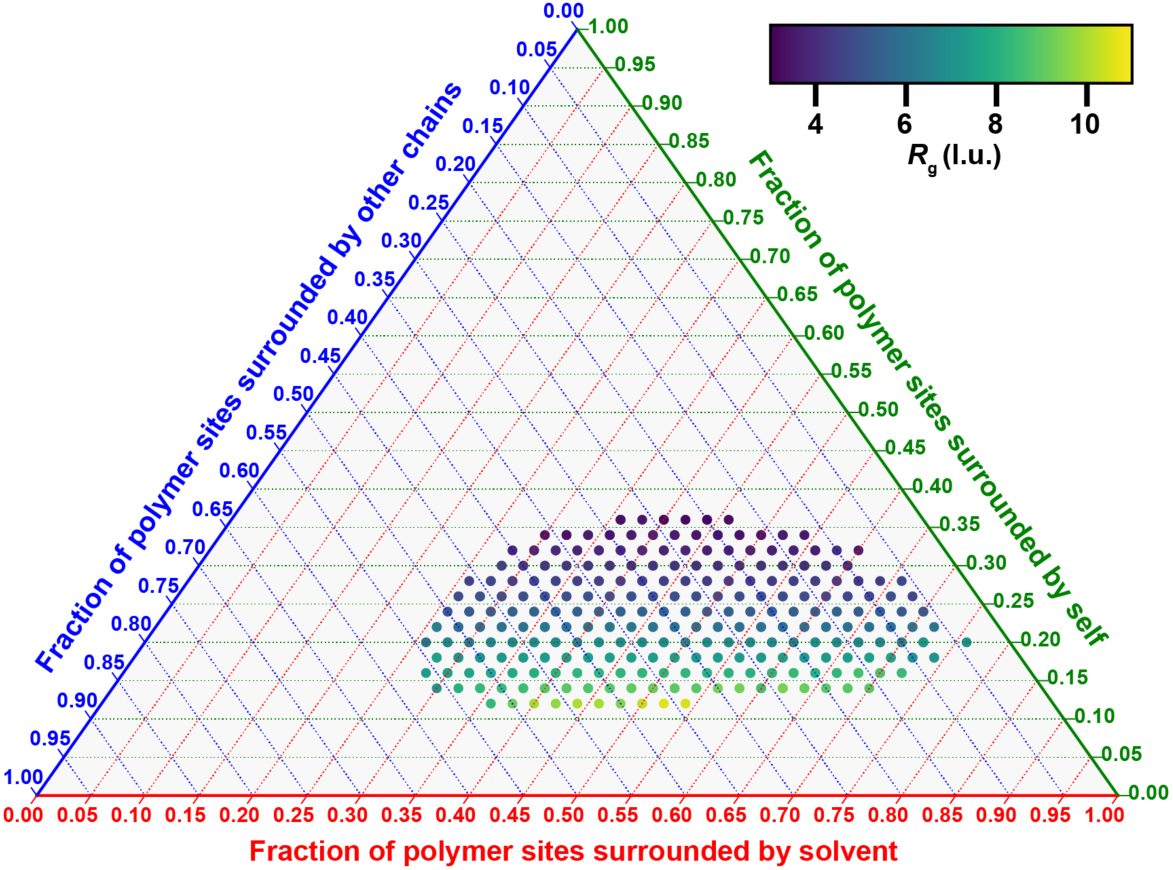
Ternary plot depicting the average radius of gyration (as represented by the color bar) of a chain in the condensate as a function of the fraction of its polymer sites that are surrounded by solvent (lower axis; red), by other chains (upper left axis; blue), or by itself (upper right axis; green). Results are shown here for the wild-type A1-LCD at ω = -4.5. The direction of the tick marks along each axis is the direction to follow for that axis. l.u. refers to lattice units.

**Extended Data Figure 6:**
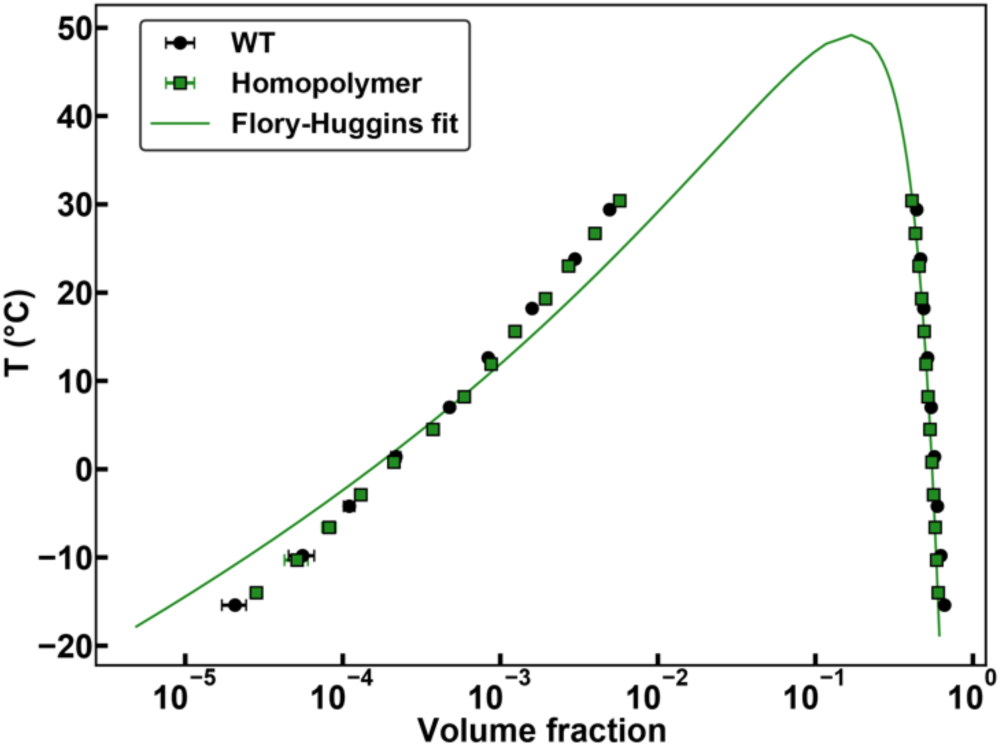
**(A)** Calculated phase diagrams of the wild-type A1-LCD and an equivalent homopolymer whose pairwise interaction energies are all set to -3.3. The solid green curve represents a fit to the Flory-Huggins theory for data obtained for the equivalent homopolymer. The method used was introduced by Martin et al., ^1^. Error bars indicate standard errors from the mean across 3 replicates. The apparent critical temperature, discerned from the fit to the mean field theory is ≈ 49°C for the solution conditions (20 mM HEPES, 150 mM NaCl, pH 7.0) that were investigated.

**Extended Data Figure 7:**
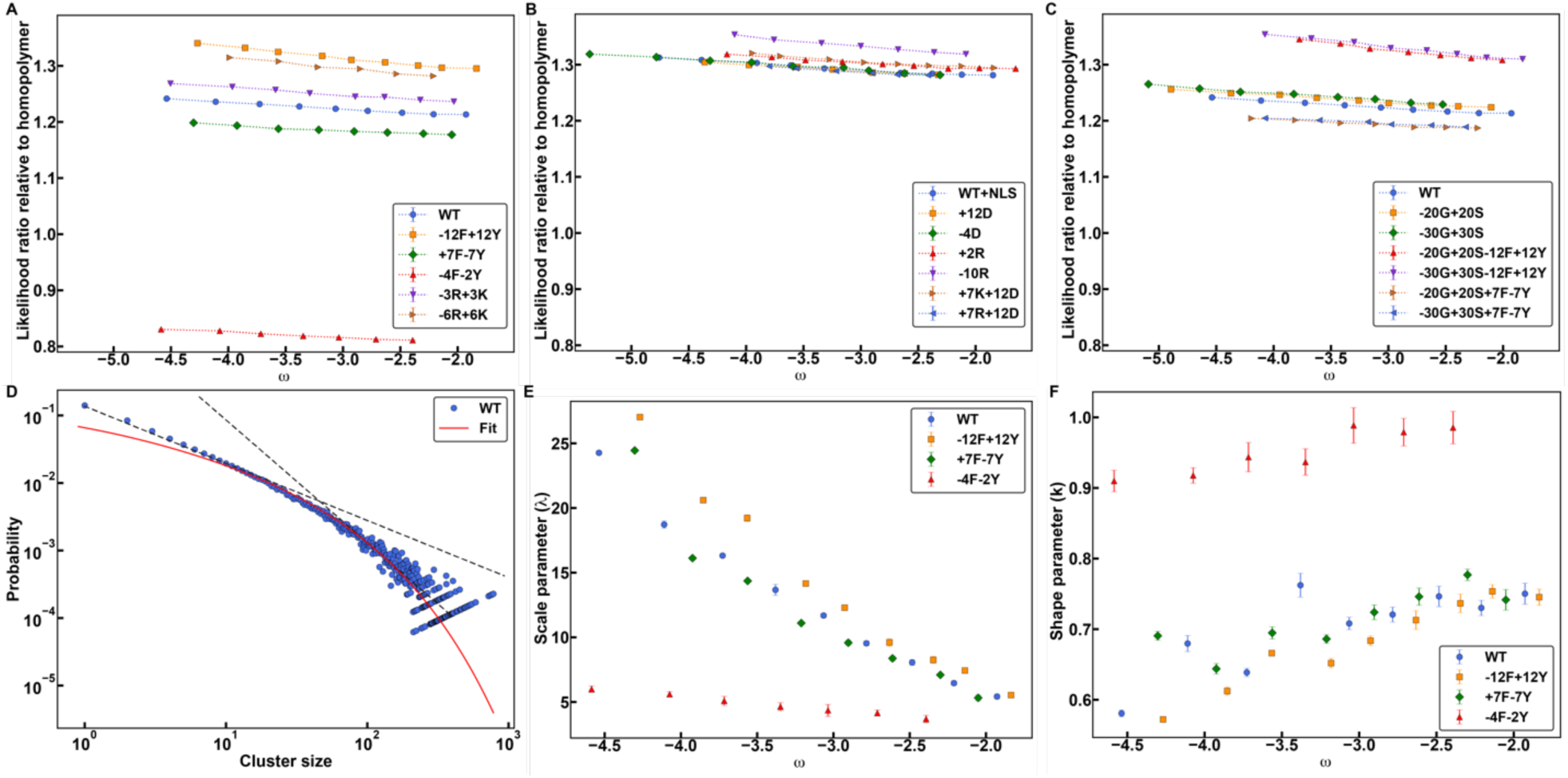
**(A-C)** Likelihood ratio that a sticker residue (Tyr or Phe) within a simulated condensate forms a crosslink with another sticker versus a spacer for variants of A1-LCD. This parameter is normalized by the respective likelihood ratio for the homopolymer, assuming the same sticker-spacer architecture as the wild-type A1-LCD. Larger values suggest that stickers within the condensate are more likely to be surrounded by other stickers. The parameter is plotted against the width of the two-phase regime, as defined in Fig. 2. The variants in **(B)** contain a nuclear localization signal (NLS), which replaces a GS motif with a PY motif. **(D)** A representative log-log plot of the probability for a sticker to be in a cluster of a particular size within the condensate for the wild-type A1-LCD at *ω* = −4.5. The cluster size is defined as the number of stickers comprising the largest connected component. The solid red curve represents a fit to the data set assuming a discrete Weibull distribution past a certain cluster size. The dashed black lines represent potential exponential fits (linear here due to the log-log scale). The poor goodness-of-fit suggests the data cannot be modeled by an exponential fit. **(E-F)** The scale parameter, λ **(E)**, and the shape parameter, k **(F)**, for the fits in **(D)** at various temperatures. Error bars in **(A-C)** indicate standard errors from the mean across 3 replicates. Error bars in **(E-F)** indicate standard deviations across 3 replicates.

**Extended Data Figure 8:**
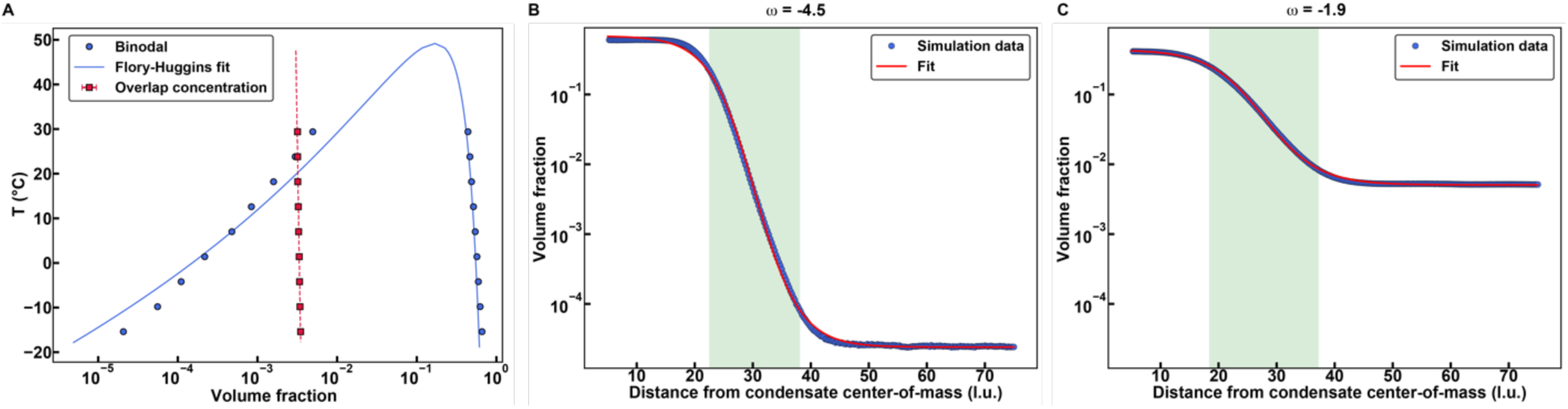
**(A)** Calculated phase diagram (circular markers) and calculated overlap concentration (square markers) of the wild-type A1-LCD. The solid blue curve represents a Flory-Huggins fit of the homopolymer phase diagram from Extended Data Figure 5 and the dashed red line represents a linear fit of the homopolymer overlap concentration to guide the eye. **(B-C)** Radial density plots of simulations of the wild-type A1-LCD at two different ω. The solid red curves correspond to fits of the data as described in the main text. The translucent green rectangles represent the interfacial regions determined by the respective fits. Error bars in **(A)** indicate standard errors from the mean across 3 replicates.

## References

1. Banani SF, Lee HO, Hyman AA, Rosen MK. Biomolecular condensates: organizers of cellular biochemistry. Nature reviews Molecular cell biology 2017, 18(5): 285–298.

2. Shin Y, Brangwynne CP. Liquid phase condensation in cell physiology and disease. Science 2017, 357(6357): eaaf4382.

3. Brangwynne CP, Eckmann CR, Courson DS, Rybarska A, Hoege C, Gharakhani J, et al. Germline P granules are liquid droplets that localize by controlled dissolution/condensation. Science 2009, 324(5935): 1729–1732.

4. Li P, Banjade S, Cheng H-C, Kim S, Chen B, Guo L, et al. Phase transitions in the assembly of multivalent signalling proteins. Nature 2012, 483(7389): 336–340.

5. Brangwynne CP, Tompa P, Pappu RV. Polymer physics of intracellular phase transitions. Nature Physics 2015, 11(11): 899–904.

6. Li P, Banjade S, Cheng HC, Kim S, Chen B, Guo L, et al. Phase transitions in the assembly of multivalent signalling proteins. Nature 2012, 483(7389): 336–340.

7. Choi J-M, Holehouse AS, Pappu RV. Physical Principles Underlying the Complex Biology of Intracellular Phase Transitions. Annual Review of Biophysics 2020, 49: 107–133.

8. Martin EW, Holehouse AS, Peran I, Farag M, Incicco JJ, Bremer A, et al. Valence and patterning of aromatic residues determine the phase behavior of prion-like domains. Science 2020, 367(6478): 694–699.

9. Bremer A, Farag M, Borcherds WM, Peran I, Martin EW, Pappu RV, et al. Deciphering how naturally occurring sequence features impact the phase behaviours of disordered prion-like domains. Nature Chemistry 2022, 14: 196–207.

10. Cohan MC, Shinn MK, Lalmansingh JM, Pappu RV. Uncovering Non-random Binary Patterns Within Sequences of Intrinsically Disordered Proteins. Journal of Molecular Biology 2022, 434(2): 167373.

11. Holehouse AS, Ginell GM, Griffith D, Böke E. Clustering of Aromatic Residues in Prion-like Domains Can Tune the Formation, State, and Organization of Biomolecular Condensates. Biochemistry 2021, 60(47): 3566–3581.

12. Bremer A, Farag M, Borcherds WM, Peran I, Martin EW, Pappu RV, et al. Deciphering how naturally occurring sequence features impact the phase behaviours of disordered prion-like domains. Nature Chemistry 2021, 14: 196–207.

13. Lantman CW, MacKnight WJ, Lundberg RD. Structural Properties of Ionomers. Annual Review of Materials Science 1989, 19(1): 295–317.

14. Cates ME, Witten TA. Chain conformation and solubility of associating polymers. Macromolecules 1986, 19(3): 732–739.

15. Rubinstein M, Dobrynin AV. Solutions of Associative Polymers. Trends in Polymer Science 1997, 5(6): 181–186.

16. Semenov AN, Rubinstein M. Thermoreversible gelation in solutions of associative polymers. 1. Statics. Macromolecules 1998, 31: 1373-1385.

17. Tanaka F. Theoretical Study of Molecular Association and Thermoreversible Gelation in Polymers. Polymer Journal 2002, 34(7): 479–509.

18. Tanaka F. Theory of Molecular Association and Thermoreversible Gelation. Molecular Gels: Materials with self-assembled fibrillar networks. Springer: Dordrecht, The Netherlands, 2006, pp 17–78.

19. Wang J, Choi JM, Holehouse AS, Lee HO, Zhang X, Jahnel M, et al. A Molecular Grammar Governing the Driving Forces for Phase Separation of Prion-like RNA Binding Proteins. Cell 2018, 174(3): 688–699 e616.

20. Choi J-M, Dar F, Pappu RV. LASSI: A lattice model for simulating phase transitions of multivalent proteins. PLOS Computational Biology 2019, 15(10): e1007028.

21. Choi JM, Hyman AA, Pappu RV. Generalized models for bond percolation transitions of associative polymers. Physical Review E 2020, 102(4-1): 042403.

22. Harmon TS, Holehouse AS, Rosen MK, Pappu RV. Intrinsically disordered linkers determine the interplay between phase separation and gelation in multivalent proteins. eLife 2017, 6: 30294.

23. Kar M, Dar F, Welsh TJ, Vogel L, Kuhnemuth R, Majumdar A, et al. Phase separating RNA binding proteins form heterogeneous distributions of clusters in subsaturated solutions. bioRxiv 2022: 10.1101/2022.1102.1103.478969.

24. Guillen-Boixet J, Kopach A, Holehouse AS, Wittmann S, Jahnel M, Schlussler R, et al. RNA-Induced Conformational Switching and Clustering of G3BP Drive Stress Granule Assembly by Condensation. Cell 2020, 181(2): 346–361 e317.

25. Franzmann TM, Jahnel M, Pozniakovsky A, Mahamid J, Holehouse AS, Nüske E, et al. Phase separation of a yeast prion protein promotes cellular fitness. Science 2018, 359(6371): eaao5654.

26. Tanaka F, Edwards SF. Viscoelastic properties of physically crosslinked networks.1. Transient network theory. Macromolecules 1992, 25(5): 1516–1523.

27. Zhang Z, Chen Q, Colby RH. Dynamics of associative polymers. Soft Matter 2018, 14(16): 2961–2977.

28. Carmesin I, Kremer K. The bond fluctuation method: a new effective algorithm for the dynamics of polymers in all spatial dimensions. Macromolecules 1988, 21(9): 2819–2823.

29. Kremer K, Binder K. Monte Carlo simulation of lattice models for macromolecules. Computer Physics Reports 1988, 7(6): 259–310.

30. Shaffer JS. Effects of chain topology on polymer dynamics: Bulk melts. The Journal of Chemical Physics 1994, 101(5): 4205–4213.

31. Ruff K, M., Harmon TS, Pappu RV. CAMELOT: A machine learning approach for coarse-grained simulations of aggregation of block-copolymeric protein sequences. The Journal of Chemical Physics 2015, 143(24): 243123.

32. Riback JA, Bowman MA, Zmyslowski AM, Knoverek CR, Jumper JM, Hinshaw JR, et al. Innovative scattering analysis shows that hydrophobic disordered proteins are expanded in water. Science 2017, 358(6360): 238–241.

33. Dignon GL, Zheng W, Best RB, Kim YC, Mittal J. Relation between single-molecule properties and phase behavior of intrinsically disordered proteins. Proceedings of the National Academy of Sciences USA 2018, 115(40): 9929–9934.

34. Zeng X, Holehouse AS, Chilkoti A, Mittag T, Pappu RV. Connecting Coil-to-Globule Transitions to Full Phase Diagrams for Intrinsically Disordered Proteins. Biophysical Journal 2020, 119(2): 402–418.

35. Hazra MK, Levy Y. Biophysics of Phase Separation of Disordered Proteins Is Governed by Balance between Short-And Long-Range Interactions. The Journal of Physical Chemistry B 2021, 125(9): 2202–2211.

36. Bremer A, Farag M, Borcherds WM, Peran I, Martin EW, Pappu RV, et al. Deciphering how naturally occurring sequence features impact the phase behaviors of disordered prion-like domains. bioRxiv 2021: 2021.2001.2001.425046.

37. Fisher ME. Shape of a Self-Avoiding Walk or Polymer Chain. The Journal of Chemical Physics 1966, 44(2): 616–622.

38. Jiang R, Murthy DNP. A study of Weibull shape parameter: Properties and significance. Reliability Engineering & System Safety 2011, 96(12): 1619–1626.

39. Newman MEJ. The structure and function of complex networks. SIAM Review 2003, 45: 167–256.

40. Erdős P, Rényi A. On random graphs. Publicationes Mathematicae 1959, 6: 290–297.

41. Patel A, Lee HO, Jawerth L, Maharana S, Jahnel M, Hein MY, et al. A liquid-to-solid phase transition of the ALS protein FUS accelerated by disease mutation. Cell 2015, 162(5): 1066–1077.

42. Jawerth L, Fischer-Friedrich E, Saha S, Wang J, Franzmann T, Zhang X, et al. Protein condensates as aging Maxwell fluids. Science 2020, 370(6522): 1317–1323.

43. Molliex A, Temirov J, Lee J, Coughlin M, Kanagaraj AP, Kim HJ, et al. Phase separation by low complexity domains promotes stress granule assembly and drives pathological fibrillization. Cell 2015, 163(1): 123–133.

44. Linsenmeier M, Hondele M, Grigolato F, Secchi E, Weis K, Arosio P. Dynamic arrest and aging of biomolecular condensates are regulated by low-complexity domains, RNA and biochemical activity. bioRxiv 2021: 2021.2002.2026.433003.

45. Shillcock JC, Lagisquet C, Alexandre J, Vuillon L, Ipsen JH. Model biomolecular condensates have heterogeneous structure quantitatively dependent on the interaction profile of their constituent macromolecules. bioRxiv 2022: 2022.2003.2025.485792.

46. Aarts DGAL, Schmidt M, Lekkerkerker HNW. Direct Visual Observation of Thermal Capillary Waves. Science 2004, 304(5672): 847–850.

47. Buff FP, Lovett RA, Stillinger FH. Interfacial Density Profile for Fluids in the Critical Region. Physical Review Letters 1965, 15(15): 621–623.

48. Bu W, Kim D, Vaknin D. Density Profiles of Liquid/Vapor Interfaces Away from Their Critical Points. The Journal of Physical Chemistry C 2014, 118(23): 12405–12409.

49. Cahn JW, Hilliard JE. Free Energy of a Nonuniform System. I. Interfacial Free Energy. The Journal of Chemical Physics 1958, 28(2): 258–267.

50. Fisk S, Widom B. Structure and Free Energy of the Interface between Fluid Phases in Equilibrium near the Critical Point. The Journal of Chemical Physics 1969, 50(8): 3219–3227.

51. Bauer BA, Patel S. Properties of water along the liquid-vapor coexistence curve via molecular dynamics simulations using the polarizable TIP4P-QDP-LJ water model. The Journal of Chemical Physics 2009, 131(8): 084709.

52. Steinhauser MO. A molecular dynamics study on universal properties of polymer chains in different solvent qualities. Part I. A review of linear chain properties. The Journal of Chemical Physics 2005, 122(9): 094901.

53. Böddeker TJ, Rosowski KA, Berchtold D, Emmanouilidis L, Han Y, Allain FHT, et al. Non-specific adhesive forces between filaments and membraneless organelles. Nature Physics 2022, 18(5): 571–578.

54. Gennes PGd. Reptation of a Polymer Chain in the Presence of Fixed Obstacles. The Journal of Chemical Physics 1971, 55(2): 572–579.

55. Ying Q, Chu B. Overlap concentration of macromolecules in solution. Macromolecules 1987, 20(2): 362–366.

56. Wei MT, Elbaum-Garfinkle S, Holehouse AS, Chen CC, Feric M, Arnold CB, et al. Phase behaviour of disordered proteins underlying low density and high permeability of liquid organelles. Nature Chemistry 2017, 9(11): 1118–1125.

57. Joseph JA, Reinhardt A, Aguirre A, Chew PY, Russell KO, Espinosa JR, et al. Physics-driven coarse-grained model for biomolecular phase separation with near-quantitative accuracy. Nature Computational Science 2021, 1(11): 732–743.

58. Tesei G, Schulze TK, Crehuet R, Lindorff-Larsen K. Accurate model of liquid-liquid phase behavior of intrinsically disordered proteins from optimization of single-chain properties. Proceedings of the National Academy of Sciences USA 2021, 118(44): e2111696118.

59. Dignon GL, Zheng W, Kim YC, Best RB, Mittal J. Sequence determinants of protein phase behavior from a coarse-grained model. PLoS Computational Biology 2018, 14(1): e1005941.

60. Amit DJ. The Ginzburg criterion-rationalized. Journal of Physics C: Solid State Physics 1974, 7(18): 3369–3377.

61. Brangwynne CP, Mitchison TJ, Hyman AA. Active liquid-like behavior of nucleoli determines their size and shape in Xenopus laevis oocytes. Proceedings of the National Academy of Sciences USA 2011, 108(11): 4334–4339.

62. Law JO, Jones CM, Stevenson T, Turner MS, Kusumaatmaja H, Grellscheid SN. Using Shape Fluctuations to Probe the Mechanics of Stress Granules. bioRxiv 2022: 2022.2005.2003.490456.

63. Bergeron-Sandoval LP, Kumar S, Heris HK, Chang CLA, Cornell CE, Keller SL, et al. Endocytic proteins with prion-like domains form viscoelastic condensates that enable membrane remodeling. Proceedings of the National Academy of Sciences USA 2021, 118(50): e2113789118.

64. Stroberg W, Schnell S. Do Cellular Condensates Accelerate Biochemical Reactions? Lessons from Microdroplet Chemistry. Biophysical Journal 2018, 115(1): 3–8.

65. Zimmermann MR, Bera SC, Meisl G, Dasadhikari S, Ghosh S, Linse S, et al. Mechanism of Secondary Nucleation at the Single Fibril Level from Direct Observations of Aβ42 Aggregation. Journal of the American Chemical Society 2021, 143(40): 16621–16629.

66. Törnquist M, Michaels TCT, Sanagavarapu K, Yang X, Meisl G, Cohen SIA, et al. Secondary nucleation in amyloid formation. Chemical Communications 2018, 54(63): 8667–8684.

67. Linsenmeier M, Faltova L, Palmiero UC, Seiffert C, Küffner AM, Pinotsi D, et al. The interface of condensates of the hnRNPA1 low complexity domain promotes formation of amyloid fibrils. bioRxiv 2022: 2022.2005.2023.493075.

68. Alshareedah I, Moosa MM, Pham M, Potoyan DA, Banerjee PR. Programmable Viscoelasticity in Protein-RNA Condensates with Disordered Sticker-Spacer Polypeptides. Nature Communications 2021, 12: 6620.

69. Zhou H-X. Viscoelasticity of biomolecular condensates conforms to the Jeffreys model. The Journal of Chemical Physics 2021, 154(4): 041103.

70. Feric M, Vaidya N, Harmon TS, Mitrea DM, Zhu L, Richardson TM, et al. Coexisting Liquid Phases Underlie Nucleolar Subcompartments. Cell 2016, 165(7): 1686–1697.

71. Meng W, Lyle N, Luan B, Raleigh DP, Pappu RV. Experiments and simulations show how long-range contacts can form in expanded unfolded proteins with negligible secondary structure. Proceedings of the National Academy of Sciences USA 2013, 110(6): 2123–2128.

72. Pedregosa F, Varoquax G, Gramfort A, Michell V, Thirion B. Scitkit-learn: Machine Learning in Python. Journal of Machine Learning Research 2011, 12: 2825–2830.

73. Newman MEJ, Strogatz SH, Watts DJ. Random graphs with arbitrary degree distributions and their applications. Physical Review E 2001, 64(2): 026118.

74. Watts DJ, Strogatz SH. Collective dynamics of ‘small-world’ networks. Nature 1998, 393(6684): 440–442.

## References

1. Martin EW, Holehouse AS, Peran I, Farag M, Incicco JJ, Bremer A, et al. Valence and patterning of aromatic residues determine the phase behavior of prion-like domains. Science 2020, 367(6478): 694–699.

